# HOX13 factors shape cell developmental transcriptional program through multiple mechanisms that regulate chromatin accessibility

**DOI:** 10.64898/2026.11.03.764042

**Authors:** Remi Robert, Rachel Paul, H. Scott Stadler, Rushikesh Sheth, Rodrigue Peraldi, Federico Gonzalez, Marie Kmita

## Abstract

HOX transcription factors pattern embryonic tissues and organs and have post-developmental roles in adult tissue maintenance and function. They can bind inaccessible chromatin, but their roles in chromatin accessibility remain unclear. Using *in vivo* and *in vitro* approaches, we show that HOX13 factors promote both initial opening and maintenance of accessibility at subsets of target regulatory elements. Mechanistically, the HOX13-dependent chromatin accessibility involves both its homodimerization and the recruitment of the SWI/SNF chromatin remodeling complex. Our findings indicate that SWI/SNF contributes to the extent of opening but is neither sufficient nor the sole determinant of HOXA13 pioneering activity. In iPSCs, HOXA13 induction rapidly reduces pluripotency gene expression while activating an early osteochondrogenic transcriptional program, which, at least in part, corresponds to direct gene regulation by HOXA13. Together, these findings identify multiple, context-dependent contributions to HOX13-mediated chromatin regulation, resulting in early lineage-associated responses.

## Introduction

The HOX family of transcription factors has crucial roles during embryonic development, controlling the establishment of the body plan, regional cell identities and organ formation^1^. Mammals have 39 *Hox* genes containing a homeobox sequence related to the Drosophila *Antp* gene, encoding sequence specific DNA binding transcription factors (TFs). These genes are conserved throughout the animal kingdom^2,3^. In mammals, *Hox* genes are grouped within the genome in four clusters (namely *HoxA, B, C* and *D*) located on distinct chromosomes, with each cluster containing 9 to 11 genes. This clustered organization is tightly linked to the developmental expression patterns of *Hox* genes both in time and space, a phenomenon referred to as spatio-temporal *Hox* collinearity^4^. The resulting sequential activation of *Hox* genes during gastrulation along the anterior-posterior axis confers cell fate and establishes distinct regional identities^5^. In addition to their essential role in patterning the main body axis, *Hox* genes are involved in the patterning of the limb^12^. Numerous genetic analyses revealed that the nested expression patterns of *HoxA* and *HoxD* genes along the anterior-posterior and proximo-distal axes define the architecture of the limb skeleton (reviewed in e.g.^6^). During limb development, *Hoxa13* and *Hoxd13* are expressed in the distal-most domain of the limb bud and the combined loss of function of *Hoxa13* and *Hoxd13* genes completely abrogates autopod (hand and foot) development^7^. In addition, these genes are involved in the development of other tissues, notably the urogenital system and the placenta^8,9,10^.

We previously showed that HOXA13 and HOXD13 factors (referred to as HOX13 hereafter) trigger digit development in part through their requirement for chromatin accessibility at distal limb-specific enhancers^11^. Such a role in promoting chromatin accessibility is consistent with results obtained in mouse ESC-derived motoneuron progenitors, where doxycycline-induced HOXC factors can bind regions of closed chromatin and promote specific anterior–posterior fate outcomes^12^. Similarly, in *Drosophila* Kc167 cells, strong HOX binding selectivity correlates with less accessible chromatin^13^ and in the developing *Drosophila* embryo, Ubx modifies chromatin accessibility, which is critical for proper appendage development^14^. For the HOX13 TFs, previous *in vivo* study could not discriminate whether HOX13 TFs mediate the switch from closed to open chromatin structure or if these factors are required for the maintenance of the accessible state, or both^11^. Moreover, the mechanism(s) by which HOX13 TFs contribute to chroman accessibility remain(s) unknown.

Homodimerization has emerged as a potential mechanism contributing to the ability of pioneer transcription factors to engage closed chromatin. For example, FOXA1 homodimerization has been associated with changes in chromatin accessibility at palindromic binding sites^15^. Similarly, studies of the bHLH pioneer factors HAND2 and NEUROD2 have suggested links between homodimer formation, nucleosomal DNA recognition and chromatin opening^16,17^. However, whether homodimerization is directly required for the pioneering activity of these factors remains unclear. Homodimerization has also been implicated in the regulatory functions of HOX transcription factors. In *Drosophila*, homodimerization of Sex combs (SCR) reduced is required for the transcriptional regulation of *fork head* and contributes to SCR homeotic function^18^. Among mammalian HOX proteins, homodimerization has been reported for HOXA1, HOXA2 and HOXA13^19,20^, although its functional significance *in vivo* remains poorly understood. Notably, the F344A mutation in the mouse HOXA13 protein impairs homodimerization and reduces HOXA13-dependent transcriptional activation of the target *EphA7* in luciferase reporter assays^21^. Together, these observations suggest that homodimerization may contribute to HOX transcriptional activity, but whether it also influences HOX-dependent chromatin accessibility is unknown.

Beyond transcription-factor dimerization, pioneer activity can involve several mechanisms that promote chromatin accessibility. Pioneer factors can recognize partially exposed DNA motifs on nucleosomes, with motif positioning, DNA-binding domain architecture and cooperative interactions influencing binding^22,23^. Some factors can also directly perturb nucleosome structure. For example, OCT4 can alter nucleosomal DNA positioning and promote cooperative binding, whereas NR5A2 can destabilize nucleosomes by competing with histone–DNA contacts and promoting DNA unwrapping^24,25^. Pioneer activity can also involve cooperation with ATP-dependent chromatin-remodeling complexes, including SWI/SNF. For example, PAX7 has been reported to promote changes in histone modifications before recruiting SWI/SNF to facilitate nucleosome displacement and enhancer activation^26^. Similarly, OCT4 cooperates with BRG1, a catalytic subunit of SWI/SNF, to support chromatin accessibility and regulatory element function^27^. Such mechanisms can depend on cellular context and chromatin state, including histone modifications, cell-cycle progression and cooperation with other transcription factor^26,28^. Importantly, pioneer-factor binding does not necessarily trigger immediate transcriptional activation but may instead establish primed or other regulatory chromatin states^29,30^.

In this work, we used a combination of *in vivo* and *in vitro* approaches to define the role of HOX13 TFs in chromatin accessibility during murine distal limb development and in undifferentiated human iPSCs. We show that HOX13 factors are required not only to trigger chromatin opening at their target regulatory elements but also to maintain the accessible state. By generating and analyzing a point mutation that alters HOXA13 capacity to homodimerize, we further show that homodimerization is required for efficient chromatin opening. Finally, using an unbiased approach to identify HOX13-associated proteins, we uncover an association with the SWI/SNF chromatin-remodeling complex and provide evidence for its recruitment to HOXA13-bound genomic regions. We further show that SWI/SNF activity contributes to HOX13-dependent chromatin accessibility and is essential for proper digit development.

## Results

### HOX13 TFs contribute to chromatin opening at HOX13 target regulatory elements

Our previous work revealed that in mouse limb buds at embryonic day 11.5 (E11.5), a subset of HOX13 target loci switch from an open to a closed chromatin state upon *Hox13* inactivation^11^. Single nuclei ATAC-seq in wild type versus *Hox13* null limb buds further identified these loci as being distal limb (presumptive digits)-specific regulatory regions, which suggested that digit formation is, at least in part, associated with HOX13-dependent chromatin accessibility^11,31^. We also found a subset of HOX13-bound loci in a closed conformation, which suggested that at these loci, HOX13 TFs can bind closed chromatin without changing accessibility. Alternatively, this could reflect a variation in the timing of chromatin opening at these HOX13 targets. To discriminate between these alternatives, we investigated the chromatin state of these targets in E12.5 distal limb buds compared to E11.5 (**Figure 1A**). Differential footprinting analysis of ATAC-seq at E11.5 and E12.5 shows that the most significant changes in chromatin accessibility are associated with the HOX13 motifs (**Figure 1B**). Accordingly, a subset of HOX13-bound loci in a closed chromatin state at E11.5 becomes open at E12.5 (**Figure 1C, Supplementary Figure 1A)**. Importantly, in *Hox13* null distal limb buds at E12.5, these sites remain in a closed chromatin state (**Figure 1C**), indicating that the opening of these sites at E12.5 are dependent on HOX13 function. Gene Ontology analysis of these newly opened sites reveals terms associated with tissue/organ development (**Figure 1D**) and *de novo* motif search uncovers the expected HOXA13 motif as the top hit (**Figure 1E**). Notably, sites becoming opened at E12.5 included HOX13-bound loci located in the regulatory landscape controlling *HoxA* and *HoxD* gene expression in distal limb buds (**Supplementary Figure 1B**). Together, these results support the hypothesis that the HOX13-dependent chromatin accessibility previously reported^11^ is associated with the capacity of HOX13 TFs to trigger the switch from closed to open chromatin.

**Figure 1:**
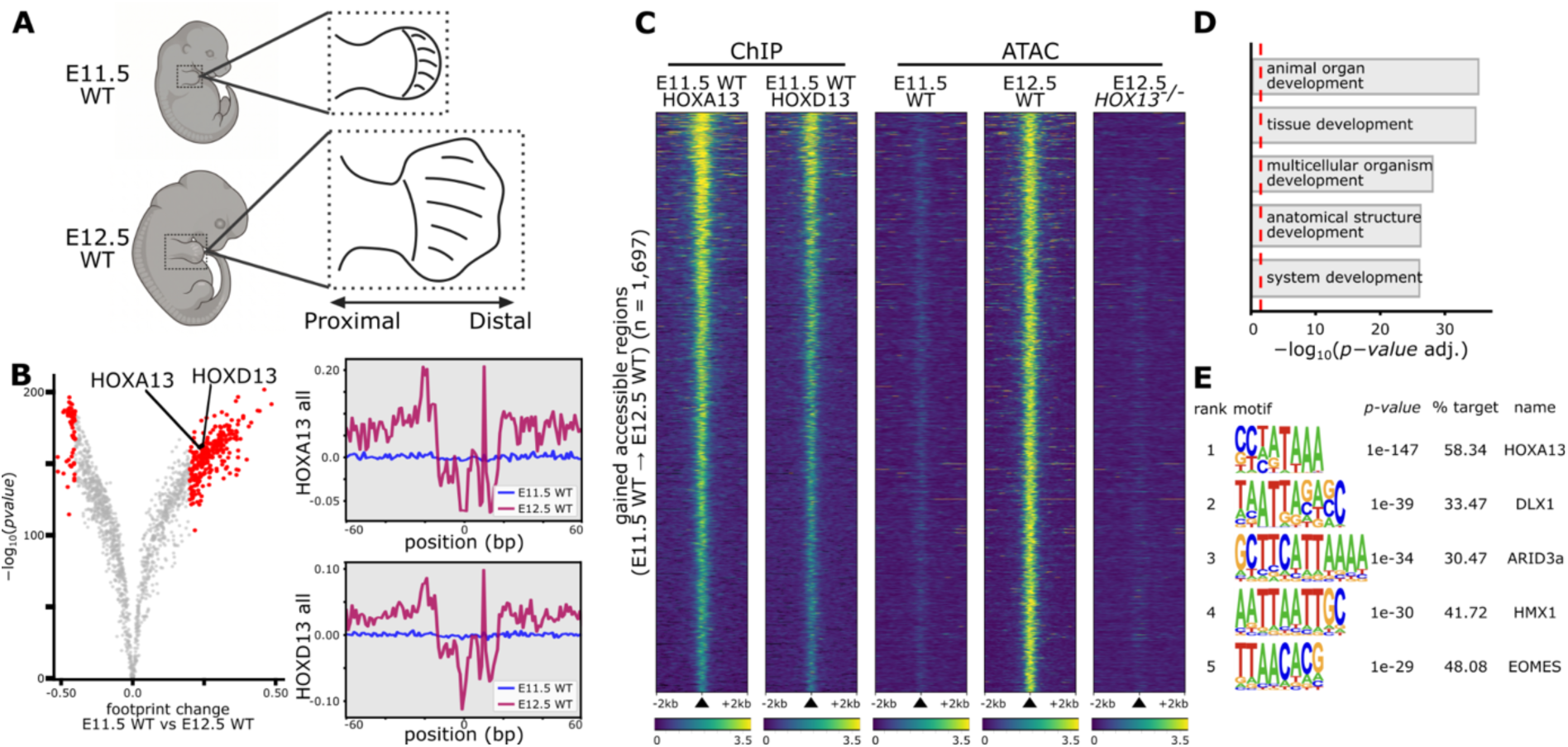
HOX13 factors remodel chromatin accessibility in mouse limb buds over time. (**A**) Schematics of the E11.5 and E12.5 mouse limb regions expressing HOX13 (striped zones) that were used for ChIP-seq and ATAC-seq. (**B**) Differential footprint analysis from E11.5 WT vs E12.5 WT ATAC-seq data (left) and aggregate footprint profiles (right) at HOXA13 and HOXD13 motifs across all sites with gained accessibility. (**C**) Heatmaps of the 1,697 gained sites at E12.5 WT bound by HOX13, showing ChIP-seq signal of HOXA13 (n = 1), HOXD13 (n = 1) at E11.5 WT and ATAC-seq signal at E11.5 WT (n = 2), E12.5 WT (n = 2) and E12.5 *Hox13 null* (n = 2) limb buds. Sites are ranked by decreasing HOXA13 ChIP-seq signal. (**D**) GREAT analysis of enriched Gene Ontology biological process terms for genes associated with the gained sites at E12.5 WT bound by HOX13 (n = 1,697). Red dashed lines indicate the cutoff (*adjusted P value* ≤ 0.05) used to define statistically significant terms. (**E**) Top scoring *de novo* motifs identified by HOMER in a 200 bp window centered on the sites for the gained sites at E12.5 WT bound by HOX13 (n = 1,697).

Because HOX13 factors are expressed in a highly dynamic and spatially restricted manner during limb development, first appearing around E10 in a small population of mesenchymal cells beneath the apical ectodermal ridge and then progressively expanding throughout the presumptive autopod over the following 48 h^32^ (**Supplementary Figure 1C-E**), the embryonic limb is not readily amenable to investigating the initial stages of HOX13 activity, when these factors may first induce chromatin opening. Notably, at E10, the scarcity of HOX13-expressing cells constrains the amount of material available for genomic analyses, while the concomitant expression of other HOX proteins may mask or confound HOX13-specific effects. We therefore established an *in vitro* system to complement our *in vivo* studies. Human induced pluripotent stem cells (iPSCs) provide a suitable HOX-negative cellular context, as they do not express *HOX* genes, can be readily expanded to generate sufficient material for genomic profiling, and can be genetically engineered to enable the controlled induction of HOX13 activity and assessment of its direct effects on chromatin accessibility and regulation. We engineered an AAVS1-targeted transgenic iPSC line with doxycycline (Dox)-inducible HOXA13 expression (hereafter iHOXA13-iPSCs; **Figure 2A**).

**Figure 2:**
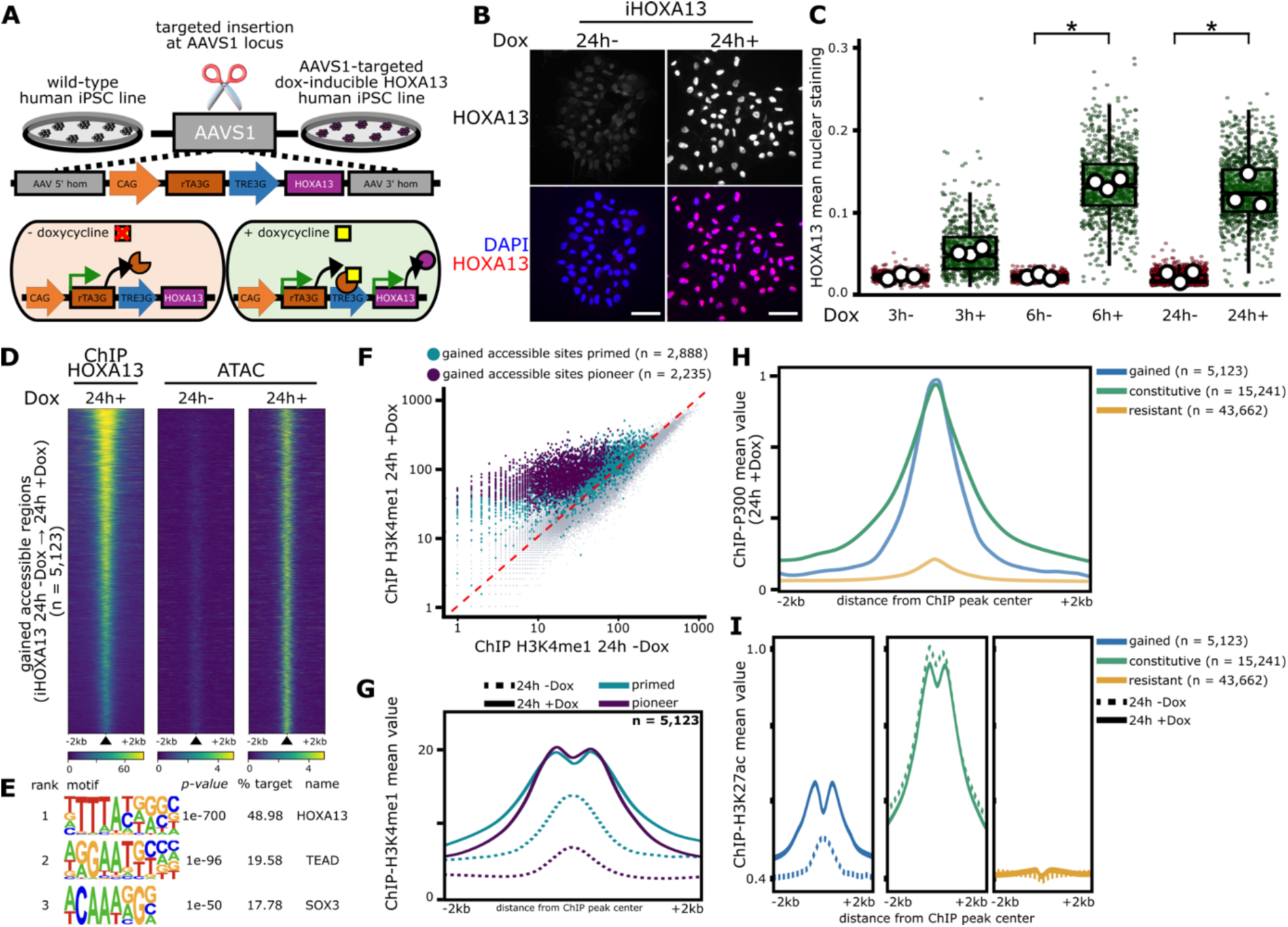
HOXA13 induction in iPSCs opens chromatin and promotes H3K4me1 and H3K27ac/P300 recruitment. (**A**) Schematic representation of the Tet-On inducible HOXA13 system (iHOXA13) integrated into the AAVS1 locus of iPSCs. (**B**) Immunostaining for HOXA13 in plated iHOXA13-iPSCs treated or not with doxycycline (Dox) during 24 h. (**C**) Boxplot of the mean nuclear HOXA13 signal intensity over time, with or without Dox (n = 3). (**D**) Heatmaps of the gained sites bound by HOXA13 (n = 5,123), showing ChIP-seq signal of HOXA13 (n = 2) at 24 h Dox treatment and ATAC-seq signal at 24 h without treatment (n = 2) or with Dox (n = 2) in iHOXA13-iPSCs. Sites are ranked by decreasing HOXA13 ChIP-seq signal. (**E**) Top scoring *de novo* motifs identified by HOMER in a 200 bp window centered on the gained sites bound by HOXA13 (n = 5,123). (**F**) Scatterplot of H3K4me1 ChIP-seq signal at ATAC-seq peaks position in iHOXA13-iPSCs after 24 h without treatment (n = 2) or with Dox (n = 2). Among the gained sites bound by HOXA13 (n = 5,123), the light blue dots indicate primed position (n = 2,888) while purple dots pioneer indicate pioneer position (n = 2,235). (**G**) Average profiles of H3K4me1 ChIP-seq signal for primed (light blue) and pioneer (purple) sites. (**H**) Average profiles of P300 ChIP-seq signal at the gained (n = 5,123; blue), constitutive (n = 15,241; green) and resistant (n = 43,662; yellow) sites bound by HOXA13, 24 h after Dox treatment (n=1) in iHOXA13-iPSCs. (**I**) Average profiles of H3K27ac ChIP-seq signal at the gained (n = 5,123; blue), constitutive (n = 15,241; green) and resistant (n = 43,662; yellow) sites bound by HOXA13, 24 h without treatment (n = 2) or with Dox (n = 2) in iHOXA13-iPSCs. For (**C)**, small dots represent individual nuclei while the larger dots represent the means of the biological replicates. Box plots: center line is the median; box limits indicate the 1st and 3rd quartiles; whiskers indicate the range. Kruskal-Wallis with non-parametric post hoc test: * if P≤0.05. Scale bars, 50 μm (**B**).

Time-course immunofluorescence analysis detected HOXA13 protein expression as early as three hours after Dox addition, reaching maximal levels by six hours (**Figure 2B-C**) and at 24h, all cells expressed HOXA13 (**Figure 2B**). Importantly, we detected no significant leaky expression of the *HOXA13* transgene in non-induced cells and no induction of endogenous *HOXA13* (**Supplementary Figure 2A**). We next examined HOXA13 genome-wide binding and its potential changes in chromatin accessibility at 24 hours of Dox treatment, allowing sufficient time for HOXA13 to exert its regulatory activity. ChIP-seq of iHOXA13-iPSCs cultured for 24 hours with or without Dox identified 73,274 loci bound by HOXA13 (**Supplementary Figure 2B-C**). ATAC-seq analysis revealed that 55,171 of these loci were inaccessible prior to HOXA13 induction, of which 5,123 became accessible following Dox treatment (**Figure 2D-E, Supplementary Figure 2D-I**). To further characterize these 5,123 HOXA13-bound regions, we investigated whether they were already marked by H3K4me1, a histone modification associated with primed or active enhancers^33^. ChIP-seq analysis at 24 hours of HOXA13 induction revealed that 2,235 of these regions lacked detectable H3K4me1 prior to Dox treatment and acquired a bimodal distribution of H3K4me1 after HOXA13 induction (**Figure 2F-G**). In contrast, the remaining 2,888 regions were already marked by H3K4me1 before Dox treatment, suggesting that they were primed prior to HOXA13 binding. Notably, at these pre-marked regions, H3K4me1 signal shifted from a centered to a bimodal distribution following induction (**Figure 2G**), consistent with nucleosome remodeling. We next asked whether these changes in chromatin accessibility and H3K4me1 distribution were accompanied by the acquisition of active enhancers features. Analysis of the entire set of 5,123 HOX13-bound sites that gained accessibility revealed P300 occupancy at levels comparable to those at constitutively open sites (**Figure 2H**). Consistent with this finding, there was a corresponding gain in H3K27ac at these sites (**Figure 2I**). Together, our *in vivo* and *in vitro* findings demonstrate that HOX13 transcription factors can engage initially closed chromatin and promote its opening, accompanied by changes in histone modifications and the acquisition of active enhancer-associated features.

### Transcriptional outcome of HOXA13 induction in iPSCs

To investigate the transcriptional consequences of HOXA13 induction, we performed RNA-seq on iHOXA13-iPSCs after 24 and 48 hours of Dox treatment. At 24 hours, 435 genes were upregulated and 387 downregulated, increasing to 1,117 and 1,599, respectively, at 48 hours (**Figure 3A, Supplementary Figure 3A-B, Supplementary Table 1**). No endogenous *HOX* genes were transcriptionally activated. Among the most strongly downregulated genes at 24 hours, were the genes encoding core pluripotency factors *NANOG*, *PRDM14* and *DNMT3b*, together with *ESRG*, *LEFTY1*, *LEFTY2*, *GDF3*, and *NODAL*, whose expression declined further at 48 hours (**Figure 3B**). Consistently, immunostaining revealed a marked reduction in NANOG and OCT4 protein levels at 24 h of HOXA13 induction (**Figure 3C**). These findings indicate that HOXA13 promotes exit from pluripotency despite the maintenance of pluripotency culture conditions. We next investigated whether this loss of pluripotency was accompanied by the activation of specific developmental programs. Gene Ontology analysis of upregulated genes at 24 h revealed enrichment for transcription regulation, cell adhesion, neuron differentiation, BMP signaling and osteoblast and chondrocyte differentiation (**Figure 3D, Supplementary Figure 3C**). Consistent with the role of HOX13 factors in limb development^7^, several genes associated with early endochondral differentiation were upregulated, including *SOX9*, a master regulator of chondrogenesis^34,35^; *BMP2* and *BMP7*, previously identified as HOX13 targets in developing limbs^36^ (**Figure 3E**). Upregulation of skeletal differentiation markers was also observed, including *RUNX2*, a key regulator of osteoblast differentiation^37,38^; and *RUNX3*, involved in chondrocyte maturation^39^. Markers of more advanced chondrogenic and osteogenic differentiation, including *COL10A1*, *IHH*, and *MMP13*, were not detected, while *SOX6*, *COMP*, *MATN1*, *GDF5*, *COL9A1*, and *CILP* were either absent or unchanged (**Supplementary Figure 3D)**. These findings raised the possibility of the activation of an early chondrogenic program without progression toward mature differentiation states at the time points examined.

**Figure 3:**
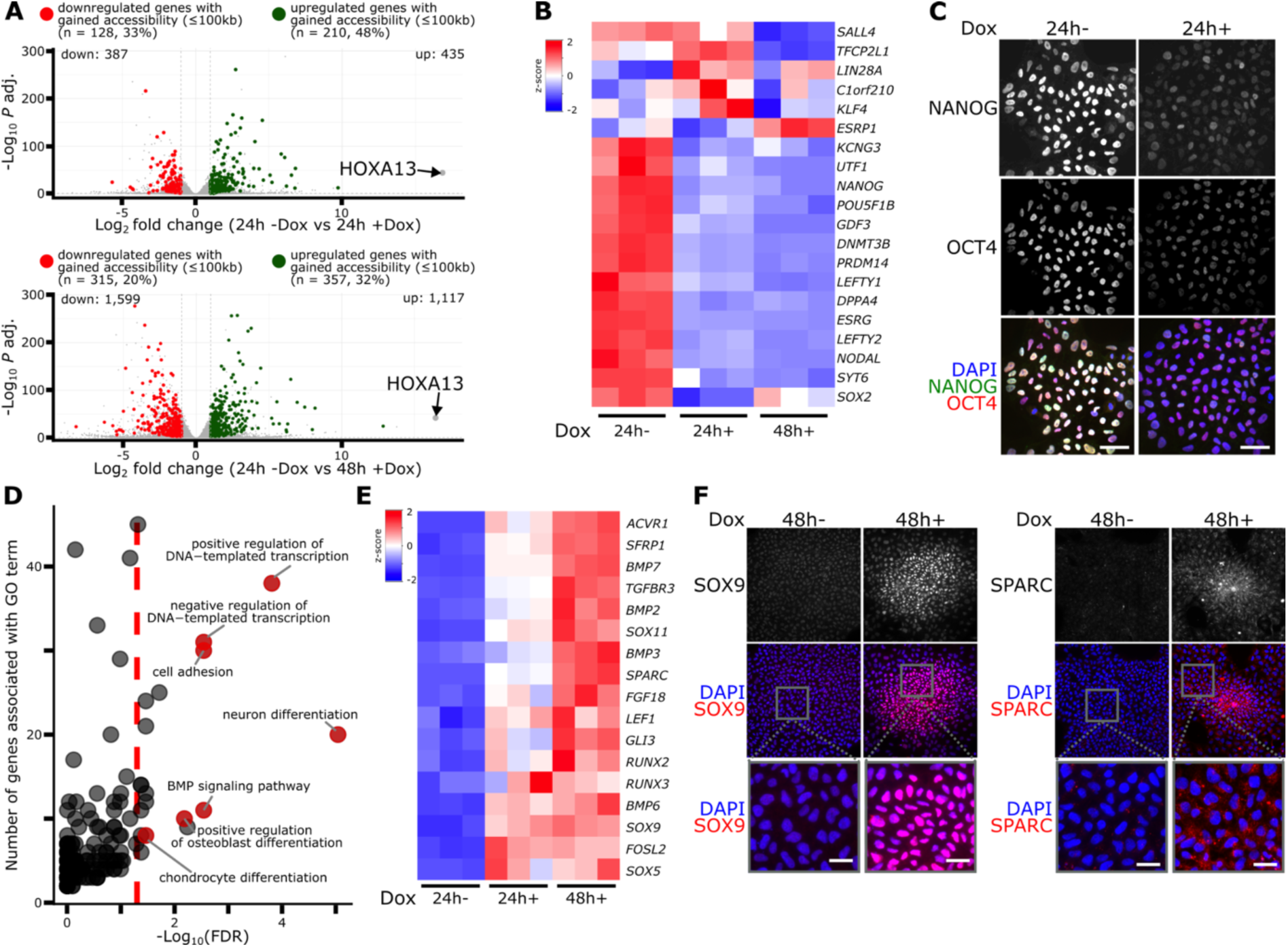
Transcriptomic analysis reveals modification in iPSCs identity upon HOXA13 induction. (**A**) Volcano plot comparing gene expression between 24h-Dox / 24h+ Dox and 24h-Dox / 48h+ Dox. The y-axis shows statistical significance (-log_10_ *P-value adjust*) and the x-axis represents log_2_ fold change. Genes with gained accessible sites within 100 kb are highlighted in red (downregulated) and green (upregulated). (**B**) Heatmap of RNA-seq gene expression (z-score) for pluripotency-associated genes in iHOXA13-iPSCs. (**C**) Immunostaining for NANOG and OCT4 in plated iHOXA13-iPSCs treated or not with Dox during 24 h. (**D**) Functional enrichment analysis (Gene Ontology: Biological Process) of the 435 genes upregulated at 24h+ Dox. The y axis shows statistical significance (-log_10_ false discovery rate, FDR) and the x-axis represents the number of genes associated with each Gene Ontology term. Red dashed lines indicate the cutoff (FDR ≤ 0.05) used to define statistically significant terms. (**E**) Heatmap of RNA-seq gene expression (z-score) for osteoblast and chondrocyte-associated genes upregulated at 24h+ Dox and/or 48h+ Dox in iHOXA13-iPSCs. (**F**) Immunostaining for SOX9 (osteochondrogenic marker) and SPARC (osteogenic mesenchymal marker) in plated iHOXA13-iPSCs treated or not with Dox during 48 h. Scale bars, 50 μm (**C**), 25 μm (**F**).

We also examined whether alternative lineage programs were activated. Despite the enrichment of neuron differentiation gene ontology terms, early neural markers (*PAX6*, *SOX1*)^41^ were not expressed, and neuronal markers (*TUBB3*, *MAP2*) were not significantly induced (**Supplementary Figure 3E**). Similarly, the absence of axial skeletal markers (*MEOX1, MEOX2, PAX1, PAX9, HES7*) and early mesodermal regulators (*EOMES, MIXL1, MSGN1, MESP1*) provided no evidence for activation of these developmental programs at the time points examined (**Supplementary Figure 3F-G**). Beyond developmental regulators, HOXA13 induction also altered the expression of genes associated with cell morphology and adhesion (**Figure 3D, Supplementary Figure 3H-I**). Notably, *MYOF* was strongly upregulated (log_2_ fold change = 4.46). Although associated with muscle development^42–44^, the absence of myogenic regulators (*PAX3, PAX7, MYOD1, MYF5, MYOG, MYMK*) argued against muscle differentiation. Instead, the concurrent upregulation of *AHNAK, ANXA1, MICAL2, EPHB3, DAAM2, ADGRG1, SPTB, ACTA1*, and *ANKRD1* suggested that the drastic upregulation of *MYOF* may be related to changes in cell organization and adhesion^45–47^.

To assess whether osteochondrogenic differentiation was occurring, we performed immunostaining for SOX9, a master regulator of chondrogenesis^34,35^, and SPARC, a matrix-regulating protein with important roles in bone formation and remodeling^40^. After 24 h of HOXA13 induction, SOX9 was detected throughout the cell population, whereas SPARC expression was heterogeneous (**Supplementary Figure 3J**). By 48h, Dox-treated cultures developed cell aggregates strongly expressing SOX9 and SPARC, while surrounding cells showed weaker or undetectable staining (**Figure 3F**). No comparable aggregation was observed in untreated controls. These morphological changes are consistent with the enrichment of cell adhesion-related gene ontology terms and, accordingly, N-Cadherin immunostaining showed a robust expression after 24h of HOXA13 induction which was not observed in the control (**Supplementary Figure 3K**). Together, these findings show that HOXA13 induction is accompanied by loss of pluripotency, activation of an early osteochondrogenic-associated transcriptional program, and changes in cell organization, without progression toward mature differentiation states at the stages examined. Whether these responses primarily reflect direct HOXA13 regulation or downstream effects remains to be established.

### HOX13 function contributes to the maintenance of chromatin accessibility

Although our data indicated that HOXA13 can trigger the switch from closed to open chromatin, it remained unclear whether this open state would persist after HOXA13 withdrawal or require its continued activity to maintain accessibility. We thus designed an experimental approach in which HOXA13 expression was transiently induced by Dox treatment for 24h, followed by an additional 24 h culture without Dox (24h+/24h-; **Figure 4A**). We first confirmed by immunostaining the clearance of HOXA13 protein following Dox withdrawal, using iHOXA13-iPSCs continuously treated with Dox for 48 h (48h+) as a control (**Figure 4B-C**). We next performed ATAC-seq to assess whether HOXA13 withdrawal affected chromatin accessibility. In iHOXA13-iPSCs continuously treated with Dox for either 24 h (24h+) or 48 h (48h+), the same set of 5,123 HOXA13-pioneered loci was accessible. In marked contrast, following Dox withdrawal (24h+/24h−), 70% of these loci (3,585 sites) reverted to a closed chromatin state, while the remaining 30% showed reduced accessibility (**Figure 4D, Supplementary Figure 4A-B**). These findings indicate that continuous HOXA13 expression is required to maintain chromatin accessibility at a large subset of HOXA13-pioneered sites.

**Figure 4:**
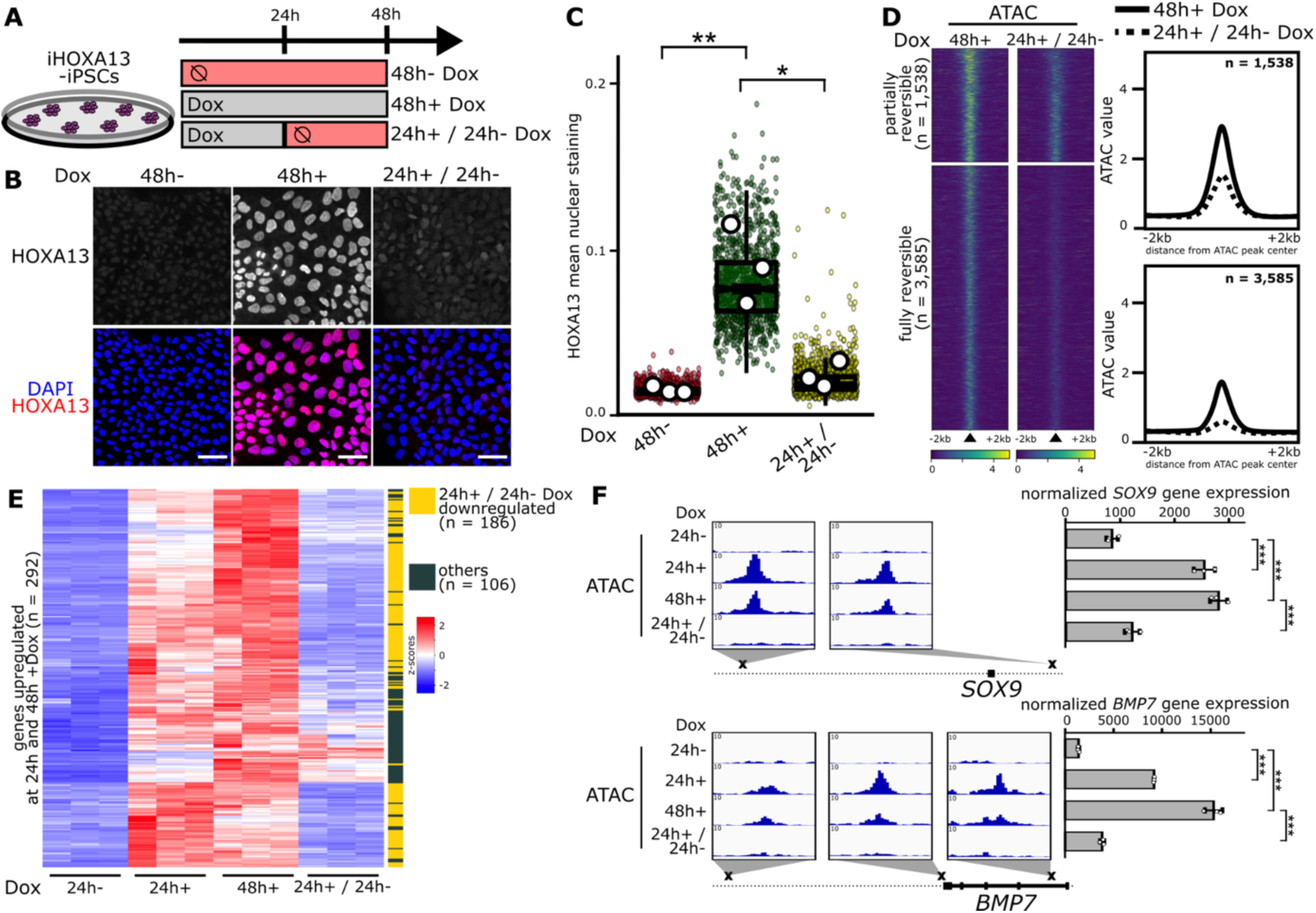
Sustained HOXA13 expression maintains chromatin accessibility and gene regulation. (**A**) Experimental approach used to profile the contribution of HOXA13 in the maintenance of chromatin accessibility. (**B**) Immunostaining for HOXA13 in plated iHOXA13-iPSCs treated or not with Dox. (**C**) Boxplot of the mean nuclear HOXA13 signal intensity in untreated cells (48h-Dox), after 48 h of Dox treatment (48h+ Dox), and after a 24 h Dox pulse followed by Dox withdrawal until 48 h (24h+/24h-) (n = 3). (**D**) Heatmaps and average profiles of the gained sites bound by HOXA13 (n = 5,123) during the first 24 h Dox treatment, showing ATAC-seq signal at 48h+ Dox (n = 2) and 24h+/24h-Dox (n = 2). Among these sites, 3,585 lost their accessibility while 1,538 decreased their accessibility. (**E**) Hierarchical clustering of RNA-seq showing transcriptional dynamic (z-score) of 292 genes up-regulated after 24 h Dox and 48 h Dox. The color bar indicates genes significantly down-regulated at 24h+/24h-Dox (yellow) and all other genes (green). (**F**) Genome browser views (IGV) of *SOX9* and *BMP7* loci, showing ATAC-seq and RNA-seq signals in 24h-Dox, 24h+ Dox, 48h+ Dox and 24h+/24h-Dox. For (**C)**, small dots represent individual nuclei while the larger dots represent the means of the biological replicates. Box plots: center line is the median; box limits indicate the 1st and 3rd quartiles; whiskers indicate the range. Kruskal-Wallis with non-parametric post hoc test: * if P≤0.05, ** if P≤0.005. Scale bars, 50 μm (**B**).

Given the increased chromatin accessibility and P300 and H3K27ac occupancy at 24 h of HOXA13 induction, we next asked whether the transcriptional responses associated with HOXA13 induction were similarly dependent on its continuous expression. To address this question, we performed RNA-seq from iHOXA13-iPSCs under the 24h+, 48h+ and 24h+/24h-conditions. Following HOXA13 withdrawal (24h+/24h-), the expression of most of the upregulated genes after 24 h and 48 h of induction, returned towards levels reminiscent of untreated cells (**Figure 4E-F, Supplementary Figure 4C-E, Supplementary Table 1**). Notably, following HOXA13 withdrawal (24h+/24h-), the expression of genes associated with early osteochondrogenic differentiation was significantly reduced or back to untreated levels (**Figure 4F**). Similarly, pluripotency marker expression shifted back toward baseline levels, resembling that of untreated cells (**Supplementary Figure 4F**).

### HOXA13 homodimerization contributes to efficient chromatin remodeling

We next investigated whether HOXA13 homodimerization contributes to its ability to promote chromatin accessibility. Homodimerization has been implicated in the pioneering activity of class I bHLH transcription factors^16^, raising the possibility that a similar mechanism may operate for HOXA13. To explore this possibility, we took advantage of the F344A mutation in mouse HOXA13 (corresponding to F346A in human HOXA13), previously shown to impair homodimerization and reduce HOXA13-dependent transcriptional activation in luciferase reporter assays^19^. Quantification of BiFC signals generated by the HOXA13^F346A^ protein revealed a twofold reduction in BiFC signal compared to wild type HOXA13 (**Figure 5A-B**), confirming the HOXA13^F346A^ mutation reduces homodimerization. We then generated an iPSC transgenic line carrying a Dox-inducible HOXA13^F346A^ transgene (referred to as iHOXA13^F346A^-iPSCs hereafter). Immunostaining performed after 24 h of Dox treatment shows that HOXA13^F346A^ was expressed in all cells (**Figure 5C-D**), reminiscent of wild type HOXA13 induced expression in iHOXA13-iPSCs (**Figure 2C, Supplementary Figure 5A**). We next investigated whether the F346A mutation affects the genomic occupancy of HOXA13. To enable quantitative comparisons of HOXA13 binding between wild-type and mutant cells, we performed spike-in-normalized ChIP-seq following Dox induction. Although HOXA13^F346A^ retained a broadly similar genome-wide binding distribution to wild-type HOXA13 (**Supplementary Figure 5B**), its occupancy was reduced at a subset of binding sites. Notably, this reduction was observed specifically at sites that gain chromatin accessibility following HOXA13 induction (**Figure 5E; Supplementary Figure 5C-F**). We next examined chromatin accessibility and for all loci opened by the wild type HOXA13, we observed a significantly reduced accessibility in iHOXA13^F346A^-iPSCs (**Figure 5E-F**), which was not the case for sites for which the accessible state was independent of HOXA13 (**Supplementary Figure 5G**). The reduced accessibility associated with the F346A mutation was more pronounced at sites where iHOXA13^F346A^ binding was decreased relative to wild-type HOXA13; however, reduced accessibility was also detected at sites where the two proteins showed equivalent binding levels (**Figure 5E-F**). We also observed gain of binding associated with the F346A mutation (**Supplementary Figure 5B**). This gain occurred at sites already open prior to HOXA13 induction and did not change chromatin accessibility. *De novo* motif search did not reveal any particular motif linked to these variations in binding efficiency (**Supplementary Figure E-F**). These results indicated that the F346A mutation modified binding affinity/stability at a small subset of its targets, all of which being in a closed chromatin state prior to HOXA13 induction and globally interfered with the efficiency of HOXA13-dependent chromatin accessibility. Consistent with these alterations, RNA-seq analysis revealed fewer differentially expressed genes following the induction of iHOXA13^F346A^ compared with wild type HOXA13 (**Figure 5G-I, Supplementary Table 2**). This included a weaker downregulation of pluripotency-associated factors and less pronounced induction of the early osteochondrogenic transcriptional program (**Figure 5J-K, Supplementary Figure 5H-I**). Together, these findings suggest that HOXA13 homodimerization influences not only genomic occupancy but also the efficiency of chromatin opening and associated transcriptional responses.

**Figure 5:**
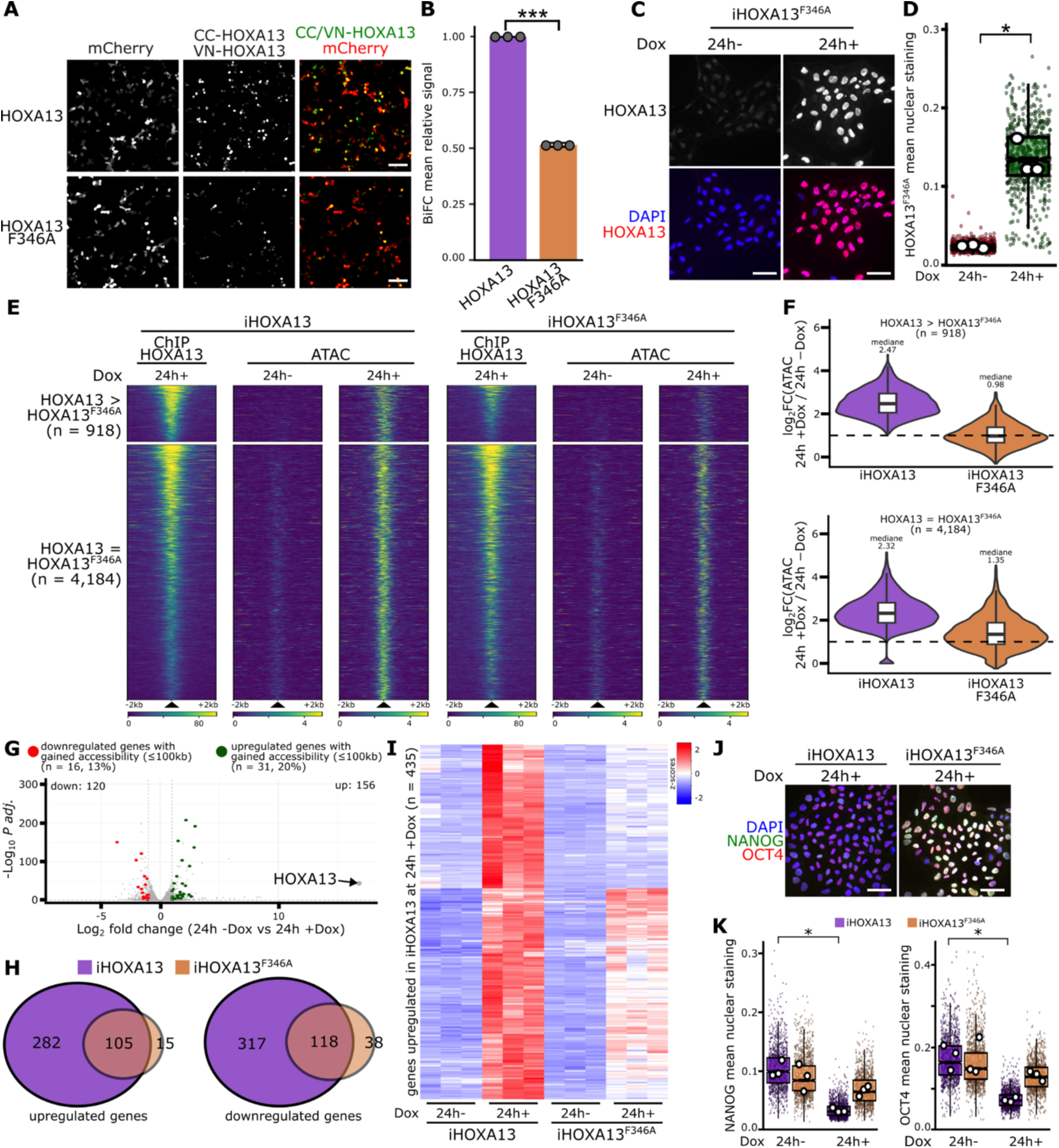
HOXA13 homodimerization is required for its recruitment to chromatin and for efficient chromatin opening. (**A**) Bimolecular fluorescence complementation (BiFC) between VN-HOXA13 and CC-HOXA13, using either wild-type or F346A mutant HOXA13, co-transfected with mCherry as a transfection control. (**B**) Quantification of the BiFC signal (mean ± S.E.M.; n = 3) for HOXA13 (purple) and HOXA13^F346A^ (orange). (**C**) Immunostaining for HOXA13 in plated iHOXA13^F346A^-iPSCs treated or not with Dox during 24 h. (**D**) Boxplot of the mean nuclear HOXA13^F346A^ signal intensity at 24 h, with or without Dox (n = 3). (**E**) Heatmaps of the gained sites bound by HOXA13 (n = 5,123) after 24 h of Dox treatment, showing HOXA13 ChIP-seq signal after 24 h of Dox and ATAC-seq signal after 24 h without treatment or with Dox in iHOXA13 and iHOXA13^F346A^-iPSCs (n = 2 for each ChIP-seq and ATAC-seq condition). Among these sites, HOXA13^F346A^ binding was decreased at 918 and unchanged at 4,184. Sites are ranked by decreasing HOXA13 ChIP-seq signal. (**F**) Violin plots with overlaid box plots showing the log_2_ fold change in ATAC-seq signal (24h+ Dox vs. 24 h −Dox) at the HOXA13 gained sites with reduced binding of HOXA13^F346A^ (n = 918) and at the gained sites with no difference in binding between HOXA13 and HOXA13^F346A^ (n = 4,184), in iHOXA13 (purple) and iHOXA13^F346A^-iPSCs (orange). (**G**) Volcano plot comparing gene expression between 24 h - Dox (n = 3) and 24h+ Dox (n = 3) in iHOXA13^F346A^-iPSC. The y-axis shows statistical significance (-log_10_ *P-value adjust*) and the x-axis represents log_2_ fold change. Among up-regulated genes, light green dots indicate genes with a gained accessible site (≤100 kb). Among down-regulated genes, light red dots indicate genes with a gained accessible site (≤100 kb). (**H**) Comparison of genes upregulated (left) and downregulated (right) upon HOXA13 induction in iHOXA13 (purple) and iHOXA13^F346A^-iPSCs (orange). (**I**) Hierarchical clustering of RNA-seq showing transcriptional dynamic (z-score) of 435 genes up-regulated after 24 h of Dox in iHOXA13-iPSCs, shown for iHOXA13 and iHOXA13^F346A^-iPSCs with or without Dox (n = 3 for each RNA-seq condition). (**J**) Immunostaining for NANOG and OCT4 in plated iHOXA13 and iHOXA13^F346A^-iPSCs treated or not with Dox during 24 h. (**K**) Boxplot of the mean nuclear NANOG and OCT4 immunofluorescence signal intensity in iHOXA13 (purple) and iHOXA13^F346A^-iPSCs (orange) after 24 h without Dox or with Dox (n = 3). For (**D)** and (**K)**, small dots represent individual nuclei while the larger dots represent the means of the biological replicates. Box plots: center line is the median; box limits indicate the 1st and 3rd quartiles; whiskers indicate the range. For (**B)**, (**D)** and **(K)**, Kruskal-Wallis with non-parametric post hoc test: * if P≤0.05, ** if P≤0.005, *** if P≤0.001. Scale bars, 100 μm (**A**), 50 μm (**C**), 50 μm (**J**).

### SWI/SNF recruitment contributes to HOX13-dependent chromatin opening

Because HOX13 TFs do not possess intrinsic nucleosome remodeling capacity, we screened developing limb tissue for potential HOX13-associated proteins using Rapid Immunoprecipitation Mass Spectrometry of Endogenous Proteins (RIME), a sensitive method for identifying endogenous protein complexes from limited starting material^48^. Among the top-ranked candidates, we recovered previously reported HOXA13 interactors, including TBX5^49^, providing independent support for the robustness of the RIME screen (**Supplementary Table 3**). Interestingly, we also found five distinct sub-units of the SWI/SNF complex, namely BRG1 (encoded by *SMARCA4*), ARID1A, SMARCC1, SMARCE1 and SMARCD2. Among these, BRG1 is one of the two alternative catalytic ATPase subunits of SWI/SNF, responsible for its ATP-dependent chromatin remodeling activity. Given the established role of SWI/SNF in ATP-dependent nucleosome remodeling, these findings raised the possibility that SWI/SNF contributes to the ability of HOX13 factors to promote chromatin opening.

To investigate the functional relevance of SWI/SNF during digit development, we first conditionally inactivated *Smarca4*, encoding BRG1, the only SWI/SNF catalytic ATPase identified in our RIME screen. We used the *Hoxa13Cre* allele, which drives recombination specifically in the presumptive autopod of developing limbs^8^. Complete loss of detectable BRG1 protein was observed from E12.5 (**Figure 6A**), approximately one day after the initiation of digit development. Conditional *Smarca4* homozygous mutants exhibited severe digit defects (**Figure 6B**), indicating that BRG1 is required for normal digit development. To determine whether BRG1 also contributes to HOX13-dependent chromatin accessibility, we performed ATAC-seq on E12.5 mutant limb buds. This analysis revealed a global reduction in accessibility at HOX13-pioneered sites following BRG1 loss (**Figures 6C-E**), supporting a role for BRG1-containing SWI/SNF complexes in establishing or maintaining accessibility at these regions. These results indicated that SWI/SNF function is critical for proper digit development.

**Figure 6:**
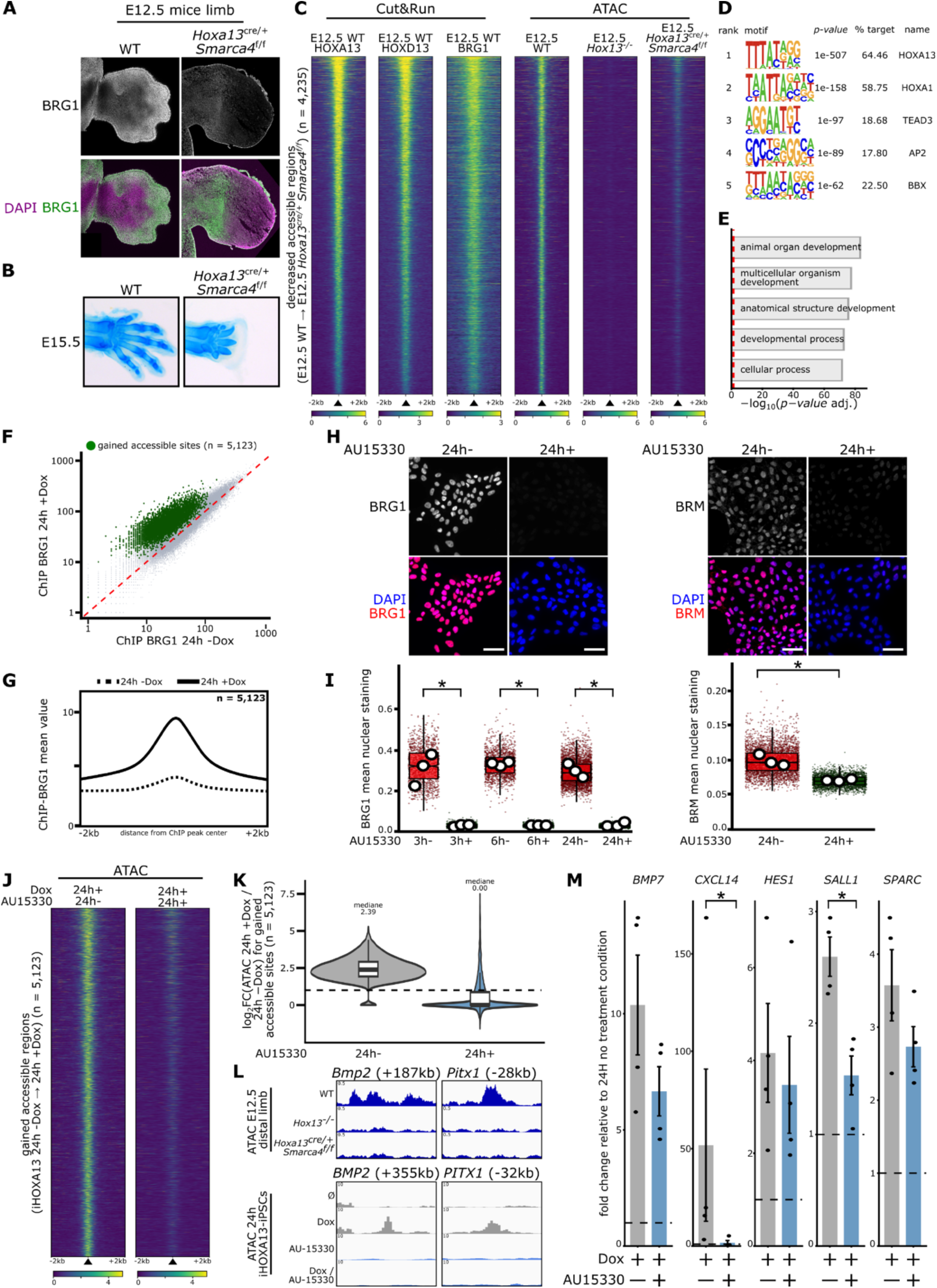
SWI/SNF recruitment is required for HOX13-mediated chromatin opening. (**A**) Immunostaining for BRG1 on sections of E12.5 limbs WT and *Hoxa13^cre/+^ Brg1^f/f^* mouse. (**B**) Skeleton of E15.5 limbs WT and *Hoxa13^cre/+^ Brg1^f/f^* mouse. The anterior is up. (**C**) Heatmaps of the sites with decreased chromatin accessibility in E12.5 Hoxa13^cre/+^ Brg1^f/f^ limb buds that are bound by HOX13 (n = 4,235), showing CUT&RUN signals for HOXA13 (n = 3) and HOXD13 (n = 3) in E12.5 WT and ATAC-seq signal in E12.5 WT (n = 2), E12.5 *Hox13* null (n = 2) and E12.5 *Hoxa13^cre/+^ Brg1^f/f^* (n = 2). Sites are ranked by decreasing HOXA13 CUT&RUN signal. (**D**) Top scoring *de novo* motifs identified by HOMER in a 200 bp window centered on the sites for the decreased sites in E12.5 *Hoxa13^cre/+^ Brg1^f/f^* bound by HOX13 (n = 4,235). (**E**) GREAT analysis of enriched Gene Ontology biological process terms for genes associated with the decreased sites in E12.5 *Hoxa13^cre/+^ Brg1^f/f^* bound by HOX13 (n = 4,235). Red dashed lines indicate the cutoff (*adjusted P value* ≤ 0.05) used to define statistically significant terms. (**F**) Scatterplot of BRG1 ChIP-seq signal at ATAC-seq peaks position in iHOXA13-iPSCs after 24 h without treatment (n = 2) or with Dox (n = 2). Sites that gain accessibility after HOXA13 binding are highlighted in green. (**G**) Average profiles of BRG1 ChIP-seq signal for sites that gain accessibility after HOXA13 binding. (**H**) Immunostaining for BRG1 and BRM in plated iHOXA13-iPSCs treated or not with AU-15330 during 24 h. (**I**) Boxplot of the mean nuclear BRG1 and BRM signal intensity with or without AU-15330 (n = 3). (**J**) Heatmaps of the gained sites bound by HOXA13 (n = 5,123) showing ATAC-seq signal after 24 h of Dox treatment, without (n = 2) or with AU-15330 (n = 2), in iHOXA13-iPSCs. Sites are ranked by decreasing HOXA13 ChIP-seq signal. (**K**) Violin plots with overlaid box plots showing the log_2_ fold change in ATAC-seq signal (24 h +Dox vs. 24 h −Dox) at the gained sites bound by HOXA13 in iHOXA13-iPSCs (n = 5,123), in the presence of AU-15330 (blue) or in its absence (grey). (**L**) Genome browser views (IGV) of *BMP2* and *PITX1* distal enhancers showing ATAC-seq signal in E11.5 mouse distal limb and in iHOXA13-iPSCs. (**M**) Real-time quantitative PCR analysis of genes upregulated by iHOXA13 (n = 4). mRNA levels were measured after 24 h of treatment with AU-15330 alone or with AU-15330 and Dox. Data are shown as fold change relative to the untreated control at the same time point. For (**I)**, small dots represent individual nuclei while the larger dots represent the means of the biological replicates. Box plots: center line is the median; box limits indicate the 1st and 3rd quartiles; whiskers indicate the range. Kruskal-Wallis with non-parametric post hoc test: * if P≤0.05. Scale bars, 200 μm (**A**), 500 μm (**B**), 50 μm (**H**).

However, because BRM, the alternative SWI/SNF catalytic ATPase, remained functional in these mutants, we could not exclude partial compensation for BRG1 loss. To further investigate the contribution of SWI/SNF to HOXA13-dependent chromatin remodeling, we therefore turned to our inducible iHOXA13-iPSC system, which allows HOXA13 activity and SWI/SNF function to be experimentally manipulated. Consistent with the involvement of SWI/SNF at HOXA13-responsive regions, we observed enrichment of ARID1A footprints at sites showing increased accessibility following HOXA13 induction (**Supplementary Figure 6A**). Furthermore, BRG1 ChIP-seq revealed increased BRG1 occupancy at HOXA13-bound sites following Dox treatment (**Figure 6F-G**). Together with the identification of SWI/SNF subunits by RIME, these results provide complementary evidence consistent with an association between HOXA13 activity and SWI/SNF occupancy at HOXA13-bound genomic regions. We next asked whether the combined depletion of BRG1 and BRM would impair HOXA13-dependent chromatin opening. To address this, we treated iHOXA13-iPSCs with the proteolysis-targeting chimera (PROTAC) AU-15330^50^, which depleted both BRG1 and BRM within 24 hours (**Figures 6H-I**). We then performed ATAC-seq following AU-15330 treatment in the presence or absence of Dox. At sites normally opened following HOXA13 induction, AU-15330 treatment reduced or abolished chromatin accessibility (**Figure 6J-K, Supplementary Figure 6B**), including at enhancers located near genes commonly regulated by HOX13 in the mouse limb and in iHOXA13-iPSCs (**Figure 6L**). These findings indicate that SWI/SNF activity contributes to the extent of HOXA13-associated chromatin opening, although they do not establish whether SWI/SNF acts during the initial opening step or subsequently contributes to remodeling or maintaining accessibility. Moreover, an increase of BRG1 occupancy was also detected at some HOXA13-bound sites that remained inaccessible (**Supplementary Figure 6C-D**), indicating that BRG1 occupancy alone is insufficient to promote chromatin opening under these conditions. Finally, qRT-PCR analysis showed that AU-15330 treatment attenuated the transcriptional response of nearby genes to HOXA13 induction (**Figure 6M**), consistent with SWI/SNF activity contributing to HOXA13-associated gene regulation.

Altogether, these results support a model in which HOXA13 activity is associated with increased SWI/SNF occupancy at HOXA13-bound genomic regions, and SWI/SNF remodeling activity contributes to efficient chromatin opening and transcriptional induction of HOXA13-responsive genes.

## Discussion

The HOX family of TFs play a central role during embryonic development by establishing positional cell fate and patterning developing embryos along the main body axes^5^. In addition to their developmental function, *Hox* are expressed and functional in post-natal and adult stages as illustrated for instance with the function of the *Hox11* genes in controlling the differentiation of *Hox-*expressing skeletal stem cells in the adult skeleton^51^. A series of *in vivo* and *in vitro* evidence points to a role of the HOX TFs in promoting chromatin accessibility^11,12,14^. *In vivo* studies of the HOX13 TFs using the developing limb as a model system showed that they are required for chromatin accessibility at regulatory elements specific to distal-limbs^11^. However, in this study, it remained unclear whether HOX13 TFs contribute to the initial opening of these regions or the maintenance of their open state, or both. Here, we provide evidence, using *in vivo* and *in vitro* approaches, that HOX13 TFs are implicated both in the initial opening and maintenance of chromatin accessibility at a subset of their target regulatory elements. In iPSCs, this is associated with the loss of pluripotency and a shift towards early osteochondrogenic-like differentiation. We further provide evidence that disrupting HOXA13 homodimerization reduces chromatin accessibility at sites with comparable HOXA13 occupancy, suggesting that homodimerization contributes to the factor’s chromatin-opening activity. Finally, we show that HOX13-dependent chromatin accessibility is, in part, associated with the recruitment of the SWI/SNF chromatin remodeler complex.

Previous studies show that some mammalian and Drosophila HOX factors can bind inaccessible chromatin, although this capacity varies among HOX proteins^12,52^. Our findings provide evidence for HOX13-mediated pioneering and maintenance of chromatin accessibility but whether these activities represent a broader property of the HOX family remains to be established. More broadly, HOX-paralog-specific pioneering activity and context-dependent regulatory networks may contribute to establishing distinct regional identities. Moreover, the involvement in maintaining chromatin accessibility may play a key role in stabilizing cell lineage in adults, for instance in the context of the adult skeleton^53,54^. Interestingly, using *in vitro* motor neuron differentiation, Bulajic et al., provided evidence that a main feature of genomic binding differences among posterior HOX factors is their differential capacities to bind closed chromatin^12^. Similarly, the study of all *Drosophila* HOX factors in Kc167 cells showed that the highest binding selectivity (in absence of co-factors) coincides with less accessible chromatin^13^. Consistent with these results, we observed that the HOXA13 point mutations impairing its ability to homodimerize (F344A in mice, F346A in humans), leads to reduced chromatin binding only at sites that are in a closed conformation prior to the induction of HOXA13^F346A^ expression. Thus, while homodimerization appears to increase HOXA13 DNA-binding only in a closed chromatin context, the latter is not the only underlying driving mechanism. Whether HOXA13 homodimerization facilitates DNA motif recognition or stabilizes protein-DNA binding remains an open question. However, because we did not identify specific DNA motifs that distinguish sites where the F346A mutation reduces HOXA13 binding, these data suggest that homodimerization may not be primarily required for DNA-motif recognition. Collectively, these data raise the possibility that distinct regional identities established by HOX factors may not rely on distinct capacities to open chromatin *per se*, but rather on differences in the binding specificity or affinity of individual HOX factors at closed chromatin. This does not preclude potential variations in pioneering activity among HOX factors in a given cell type, as suggested by the comparison of HOXC9 and HOXC10 during *in vitro* motor neuron differentiation^12^. Alternatively, co-factors could be involved in differential pioneering activity of the various HOX factors at closed chromatin.

HOXA13 induction in iPSCs has a drastic impact on cell identity, with the robust down-regulation of genes encoding for the core pluripotency markers, even though the iHOXA13-iPSCs were cultured in a media favoring pluripotency. At the stages examined, this exit of the pluripotent state is accompanied by a transition towards an early osteochondrogenic specification, with, in particular, the robust upregulation of *SOX9*, a master regulator of chondrogenesis^34,35^, *BMP2*, which helps drive cartilage formation, notably in developing limbs^55^ and *RUNX2,* essential for skeletal development^37,38^. Interestingly, at 48 h of HOXA13 induction, iHOXA13-iPSCs start forming cell aggregates. Consistent with this observation, gene ontology analysis of upregulated genes highlighted cell adhesion among the top processes. Aggregation of iHOXA13-iPSCs mirrors the reassociation of dissociated HOXA13-expressing mesenchymal cells from chick or mouse limb buds, and their subsequent segregation from HOXA13-negative cells^56–58^. The iHOXA13-iPSCs cell clustering is associated with robust SOX9 and SPARC expression, while surrounding cells do not express these markers, consistent with an early chondrogenic or mesenchymal-condensation response. Although early osteochondrogenic markers typically emerge over several weeks in conventional iPSCs differentiation protocols^59–61^, HOXA13 induction activated genes associated with early osteochondrogenesis within 24 h, even under pluripotency-supporting conditions. This response included BMP2 and BMP7, previously identified as direct HOXA13 targets during limb development^36^, suggesting that HOXA13 can rapidly engage components of its developmental transcriptional program outside its normal cellular and developmental contexts. Collectively, our data indicate that HOXA13 induction in iPSCs rapidly activates a broad osteochondrogenic-associated transcriptional program, including both chondrogenic-and osteogenic-associated regulators. This 24-hour response is rapid relative to published pluripotent-cell protocols that generate chondrogenic or osteogenic cells over days to weeks ^59–61^, however, early marker induction does not by itself establish lineage commitment, and further investigation is warranted to resolve this question. Whether under more permissive differentiation conditions, these cells would progress toward a lineage specified by HOXA13 during embryonic development, remains an open question.

Our data provide evidence that inducing HOXA13 expression in iPSCs results in the recruitment of the SWI/SNF complex at HOXA13-bound sites. However, this recruitment is not sufficient to trigger the switch from closed to open chromatin as upon SWI/SNF inhibition, many HOXA13-bound sites still show ATAC peaks, though reduced compared to controls. Moreover, we also observed SWI/SNF recruitment at HOXA13-bound loci that remain in a closed conformation (resistant sites) further supporting that SWI/SNF function is not sufficient to initiate chromatin opening. Consistent with these results, *in vitro* functional analysis of the pioneer factors PU.1 and C/EBPα/β revealed that SWI/SNF is not the primary determinant of initial opening^62^. Similarly, analysis of the PAX7 pioneering activity in a mouse pituitary tumor cell line (AtT20) have provided evidence that early chromatin changes involve histone-mark modifiers, while SWI/SNF is recruited later, after cell division, to displace nucleosomes and activate enhancers^26^. In this latter study, the authors identify the deposition of the activating H3K4me1 mark as part of the first step of Pax7 pioneer action^26^. Upon HOXA13 induction in iPSCs, a subset of HOXA13-pioneered sites showed increased H3K4me1 signal. At the other subset, H3K4me1 was already present but its distribution became bimodal following HOXA13 induction, a pattern consistent with local nucleosome remodeling. These findings suggest that HOXA13-associated chromatin opening involves changes in H3K4me1 distribution, although the events that initiate opening remain to be defined.

Collectively, our findings provide evidence that multiple mechanisms shape HOXA13-mediated chromatin accessibility, although the mechanism underlying initial opening remains to be identified. Interestingly, in iPSCs, HOXA13 induction is accompanied by a rapid loss of pluripotency and activation of gene expression programs associated with early osteochondrogenic differentiation, consistent with the role of HOX13 factors in limb skeletal development. A compelling direction for future research is to determine whether more permissive differentiation conditions can promote progression of these cells toward an embryonic lineage associated with HOX13 and to investigate how cellular context shapes cell-fate specification by HOX13 and other HOX factors.

## Methods

### Mouse lines

*Hoxa13* and *Hoxd13* lines were previously reported^7^. To generate the conditional inactivation of *Smarca4* in the developing autopod, the *Smarca4* floxed line (kindly provided by Dr Julie Lessard) was crossed with the *Hoxa13:cre* line^8^. All mice were maintained in a mixed background (C57BL/6× 129). The mice were housed in a 12 h light: dark cycle between 18 and 26 °C temperature with relative humidity of 30-70% and given ad libitum access to food and water for the duration of the study. Noon of the day of the vaginal plug was considered as E0.5. Mice and embryos were genotyped by polymerase chain reaction (PCR) using genomic DNA extracted from tail biopsy specimens and yolk sacs, respectively.

### hiPSC culture

All experiments were performed using the KOLF2.1J human induced pluripotent stem cell (iPSC) line^63^ (JAX iPSC Catalog JIPSC001000). The integrity of the iPSCs was validated by assessing copy number variations with iCS-digital™ PSC test (Stem Genomics) and by karyotyping. KOLF2.1J cells were maintained at 37 °C with 5% CO2 on vitronectin (VTN-N)-coated plates (Gibco, A400457) in StemFlex medium (Gibco, A3349401) supplemented with 1X Penicillin Streptomycin (ThermoFisher, 15140122) and amplified using Versene solution (Gibco, 15040066) clump-based passaging. Before freezing in Cryostor CS10 (Stem Cell Technologies, 100-1061) each new batch of iPSCs, the culture medium was tested for mycoplasma contamination using the PCR Mycoplasma Detection Set (Takara, 6601). No contamination was detected during the study. iPSCs were thawed in presence of Y-27632 (Stem Cell Technologies, 72304) and the culture medium was changed every day. The molecules and concentrations used in this study are listed in Supplementary Table 4

### Generation of iHOXA13 and iHOXA13^F346A^ transgenic hiPSC lines

The pAAVS1-iHOXA13 and pAAVS1-iHOXA13^F346A^ donor constructs were generated through two sequential cloning steps. First, the LoxP-SA-T2A-Puro-bGHpA cassette of pAAVS1-PDi-CRISPRn (Addgene, #73500; Mandegar et al., Cell Stem Cell, 2016) was replaced with an FRT-flanked puromycin gene-trap cassette (FRT-SA-T2A-Puro-bGHpA-FRT). Subsequently, the CRISPRn cassette was replaced with a synthetic coding sequence (CDS) encoding either wild-type human HOXA13 or its F346A mutant (GenScript). For targeted integration, 2 million of iPSCs were single cell dissociated using TripleExpress (Gibco,12604013), resuspended in 200 µL of nucleofection mixture containing 1 µg AAVS1-TALEN-L, 1 µg AAVS1-TALEN-R^64^ (Addgene, #59025 and #59026 respectively), and 8 µg of either pAAVS1-iHOXA13 or pAAVS1-iHOXA13^F346A^ donor plasmid. Nucleofection was performed using an Amaxa Nucleofector II device and the Human Stem Cell Nucleofector Kit 1 (Lonza, VPH-5012), using program B-016. Following nucleofection, cells were pooled and seeded onto two VTN-N-coated 10-cm culture dishes (Sarstedt, 83.3902) in StemFlex medium (Gibco, A3349401) complemented with CloneR™2 (StemCell Technology, 100-0691). Next day, cells were selected using 0.15 µg/mL puromycin (Gibco, A1113803) for three days, followed by 0.3 µg/mL for an additional three days. After antibiotic selection, 24 resistant colonies were selected based on morphology, mechanically dissociated, and transferred individually into 24-well plates (Sarstedt, 83.3922). Colonies were expanded to near confluence, passaged using Versene (Gibco, 15040066), and replica-plated into six-well plates (Sarstedt, 83.3920). Once confluent, one replicate was used for genomic DNA extraction and Southern blot analysis, while the other was cryopreserved.

### Genotyping and quality control of iHOXA13 and iHOXA13^F346A^ transgenic hiPSC lines

Southern blot analysis was performed as previously described^64^. Briefly, genomic DNA was digested with NcoI-HF (NEB, R3193L). Hybridization with a 5′ internal probe (INT) detected fragments corresponding to the wild-type allele (5,992 bp), correctly targeted iHOXA13 or iHOXA13^F346A^ alleles (4,168 bp), and randomly integrated transgenes (variable fragment sizes). Hybridization with a 3′ external probe (EXT) identified fragments of 5,992 bp and 4,482 bp corresponding to the wild-type and correctly targeted alleles, respectively. Clones carrying a heterozygous targeted integration were selected for further quality control. Inducible expression of HOXA13 and HOXA13^F346A^-iPSCs was assessed in three independently targeted clones per line by immunostaining with an anti-HOXA13 antibody (rabbit serum, provided by the Stadler lab) after culture with or without doxycycline (Dox) (Fisher Scientific, BP26535). Based on these quality control analyses, clones iHOXA13 #2 and iHOXA13^F346A^ #8 were selected for subsequent experiments.

### ChIP and CUT&RUN samples preparation for sequencing

For mice limb, Hoxa13/d13 ChIP-seq data shown in Figure 1.C were previously published^65^ in accession number GSM2151013 and GSM2151014. Hoxa13/d13 CUT&RUN data shown in Figure 6.C were previously published^66^ in accession number GSE194114.

For iHOXA13 and iHOXA13^F346A^ iPSC, 10 million cells (for transcription factors) or 6 million cells (for histone marks) were seeded across three VTN-N-coated 10-cm culture dishes (Sarstedt, 83.3902) in StemFlex medium (Gibco, A3349401) supplemented with 10 µM Y-27632 (Stem Cell Technologies, 72304). The next day, cells were treated with or without 2 µg/mL doxycycline (Fisher Scientific, BP26535). At the end of the treatment, cells were washed three times with cold phosphate-buffered saline (PBS), scraped and pooled in a 1.5-mL tube. Chromatin was cross-linked differently depending on the antibody used: for HOXA13 and BRG1, cells were incubated with 2 mM disuccinimidyl glutarate (ThermoFisher, H58208.MC) for 45 min at room temperature (RT), followed by 1% formaldehyde (Sigma-Aldrich, F8775) for 13 min at RT; for H3K4me1 and H3K27ac, with 1% formaldehyde for 13 min at RT; and for p300, with 1% formaldehyde for 10 min at 37 °C. After quenching the cross-linking with 125 mM glycine for 10 min at RT, cells were sequentially incubated in buffer A (0.25% Triton, 10 mM Tris pH8, 10 mM EDTA, 0.5 mM EGTA) for 10 min at 4 °C and in buffer B (200 mM NaCl, 10 mM Tris pH8, 10 mM EDTA, 0.5 mM EGTA) for 30 min at 4 °C. Samples were then sonicated on a Covaris E220 in buffer CD (0.5% sodium dodecyl sulfate (SDS), 0.5% Triton, 0.05% NaDOC, 10 mM Tris pH8, 140 mM NaCl, 1 mM EDTA, 0.5 mM EGTA) for HOXA13, BRG1, H3K4me1 and H3K27ac, or low salt (1% Triton, 0,1% SDS, 150 mM NaCl, 20 mM Tris pH8, 2 mM EDTA) for p300. All buffers were supplemented with protease inhibitor cocktail and PMSF. Protein A and Protein G Dynabeads (Thermofisher, 10001D and 10003D) were incubated for 6 h at 4 °C with primary antibody. Chromatin from iPSCs was supplemented with 2-5% spike-in chromatin from mouse limb before being coupled to the beads overnight at 4 °C. The immunoprecipitated samples were then sequentially washed in low salt (1% Triton, 0,1% SDS, 150 mM NaCl, 20 mM Tris pH8, 2 mM EDTA), high salt (1% Triton, 0,1% SDS, 500 mM NaCl, 20 mM Tris pH8, 2 mM EDTA), LiCl (1% NP-40, 250 mM LiCl, 10 mM Tris pH8, 1 mM EDTA) and TE buffer (50 mM NaCl, 10 mM Tris pH8, 1 mM EDTA). The DNA was then purified on QIAquick columns (Qiagen, 28106). Library and flow cells were prepared by the IRCM Molecular Biology Core Facility according to Illumina’s recommendations and sequenced on an Illumina NovaSeq instrument with a 100 or 150-cycle paired-end configuration. Primary antibodies are listed in Supplementary Table 4.

### ATAC samples preparation for sequencing

Distal forelimb buds from wild-type embryos at E11.5 and E12.5, and from *Hox13*−/− embryos at E12.5, were dissected in cold PBS. Dissected tissue was incubated in a dissociation buffer (450 µL of 0.25% trypsin-EDTA, 50 µL of 10% bovine serum albumin (BSA) and 1 µL of DNase I) for 10 min at 37 °C. Cells were dissociated by gently pipetting up and down 10-15 times, and fetal bovine serum was then added to a final concentration of 10%. The suspension was filtered through a 40-µm nylon cell strainer and centrifuged at 300 *g* for 5 min at 4 °C. The pellet was resuspended in 1× PBS with 0.1% BSA, and 50,000 cells were counted and processed as described below for iPSCs.

In iHOXA13 and iHOXA13-F346A-iPSCs, cells were dissociated with TrypLE Express (Gibco, 12604013) and 50,000 single cells were counted. Cells were washed in PBS and incubated on ice for 30 min in hypotonic lysis buffer (0.1% (w/v) sodium citrate tribasic dihydrate, 0.1% (v/v) Triton), then in lysis buffer (10 mM Tris-HCl pH 7.4, 10 mM NaCl, 3 mM MgCl2, 0.1% (v/v) IGEPAL CA-630) for 30 min on ice. After centrifugation the resulting pellet of nuclei was resuspended in transposase Master Mix (1.25 µl 10× TD buffer, 5 µl H_2_O and 6.5 µl of Tn5: Illumina Nextera Kit; FC-121-1031) and incubated for 30 min at 37 °C. Samples were purified using MinElute PCR purification column (Qiagen, 28004). The DNA libraries and sequencing flow cells were prepared by the IRCM Molecular Biology Core Facility on an Illumina NovaSeq with 100 bp paired-end reads, according to Illumina’s recommendation.

### RNA samples preparation for sequencing

RNA was extracted from iHOXA13 and iHOXA13^F346A^ iPSCs using RNAeasy Plus mini kit (Qiagen, 74134). Ribosomal depletion, library preparation and flow-cell preparation for sequencing were performed by the IRCM Molecular Biology Core Facility according to Illumina’s recommendations. Sequencing was done on an Illumina NovaSeq instrument with a 100-cycle paired-end configuration.

### ChIP-seq and ATAC-seq data analysis

After quality control of FASTQ files with FastQC v0.12.1, ChIP-seq and ATAC-seq paired-end reads were trimmed with fastp v0.24.0 and aligned with Bowtie 2 v2.5.4 (--very-sensitive preset) to the mm10 or hg38 genome. For samples with mouse spike-in chromatin, reads were aligned to a combined hg38 / mm10 reference genome. Reads overlapping blacklisted regions were removed with bedtools v2.31.0, duplicate reads were removed with Picard v3.1.0, and sorted BAM files were generated with samtools v1.22.1. For spike-in normalization of ChIP-seq, the number of reads mapping to the mouse spike-in chromatin (mm10) was determined for each iPSCs sample, and sample-specific scaling factors were calculated as *S*ᵢ = *T* / *N*ᵢ, where *T* is a fixed target of 10⁶ spike-in reads and *N*ᵢ is the spike-in read count of sample *i*. Peaks were called with MACS3 v3.0.3 (*P-value* ≤ 1 × 10⁻⁵) using input control. Peaks detected in at least one of the two replicates of a condition were retained using the mergePeaks function of HOMER v4.11, and the peak lists from the two conditions were pooled to obtain the set of peaks used for differential analysis. Read counts were quantified with HOMER v4.11, scaled with the sample-specific scaling factors, and differential enrichment was assessed with DESeq2. Peaks with |log2 fold change| ≥ 1 and adjusted *P-value* ≤ 0.05 were considered significant. Among these, peaks already detected in both conditions were classified as increased or decreased, whereas peaks detected in only one condition were classified as gained or lost. ChIP-seq peaks were annotated with ChIPseeker v1.8.6, and GenomicRanges v1.24.1 was used to associate peaks with genes within a 100-kb window. GREAT v2.14.0 was used to identify biological processes enriched among genes associated with the ChIP-seq peaks. ATAC-seq footprinting analysis was performed with TOBIAS v0.16.1. *De novo* motif analysis of ChIP-seq peaks was performed with the findMotifsGenome.pl command of HOMER v4.11. For visualization, bigWig files were generated with HOMER v4.11. BigWig files from replicates of the same condition were merged, and heatmaps with average profiles were generated with deepTools v3.5.6.

### RNA-seq data analysis

After quality control of the FASTQ files with FastQC v0.12.1, paired-end reads were aligned to the hg38 reference genome with STAR v2.7.11 using default parameters. Reads were counted with featureCounts v2.1.1 using default parameters, with reverse-stranded library settings. Differential expression analysis was performed in R (RStudio) with DESeq2 v1.52.0 using default parameters on the count matrix. Only genes with DESeq2-normalized counts ≥ 100 were retained, and genes with |log2 fold change| ≥ 1 and adjusted *P-value* ≤ 0.05 were considered differentially expressed. Functional annotation and enrichment analysis of the differentially expressed genes were performed with DAVID. The RNA-seq data are provided in Supplementary Table 1 and 2.

### RIME and analysis

Rapid Immunoprecipitation Mass Spectrometry (RIME) experiments were performed as described^67^. E11.5 forelimb buds were fixed 45 minutes with DSG (ThermoFisher, H58208.MC) at RT and then 13 minutes with 1% formaldehyde (Sigma-Aldrich, F8775) at RT. The pellet of fixed chromatin was snap-frozen and kept at −80 °C. Briefly, fixed chromatin was incubated 10 minutes in LB1 (50 mM HEPES-KOH pH 7.5, 140 mM NaCl, 1mM EDTA, 10% (v/v) glycerol, 0.5% (v/v) NP-40, 0.25% (v/v) Triton) on ice, then 5 minutes in LB2 (10 mM Tris-HCl pH8.0, 200 mM NaCl, 1 mM EDTA, 0.5 mM EGTA) on ice, and resuspended in LB3 buffer (10 mM Tris-HCl pH8.0, 100 mM NaCl, 1 mM EDTA, 0.5 mM EGTA, 0.1% (w/v) sodium deoxycholate, 0.5% (v/v) N-Lauroylsarcosine) (all buffers were supplemented with protease inhibitor cocktail and PMSF). Chromatin was then sonicated with a Covaris E220. After sonication, Triton was added (1% final) and the lysate was centrifuged for 10 minutes at 20 000g at 4 °C. The lysate was then resuspended with an antibody coupled to magnetic beads and incubated overnight at 4 °C. Beads were washed ten times with RIPA buffer (50 mM HEPES pH 7.6, 1 mM EDTA, 0.7% (w/v) sodium deoxycholate, 1% (v/v) NP-40, 0.5 M LiCl), then four times with 50 mM ammonium bicarbonate (AMBIC), and finally resuspended in 50 mM AMBIC. Following this step, the final steps of enzymatic digestion and mass spectrometry analysis were performed by the Proteomic platform at IRCM. RIME assays were performed on six biological replicates, each using 40–50 limb buds. Proteins detected in at least four of the six replicates were considered putative HOXA13-interacting proteins. Immunoprecipitations were carried out with either 1 µg of rabbit IgG (Millipore) as a control or 5 µg of an affinity-purified rabbit anti-HOXA13 antibody (provided by the Stadler lab). Antibodies were coupled to a mix of Protein A and Protein G magnetic beads (Thermo Fisher, 10001D and 10003D) and incubated for 6 h at 4 °C. The RIME data are provided in Supplementary Table 3.

### Bimolecular fluorescence complementation and quantification

HEK-293 cells were cultured in Dulbecco’s modified Eagle’s medium (DMEM GlutaMAX-I; Gibco) supplemented with 10% (v/v) heat-inactivated fetal bovine serum, 1% (v/v) penicillin-streptomycin (5,000 U/mL penicillin and 5 mg/mL streptomycin) and 1 mM sodium pyruvate, at 37 °C with 5% CO2. For BiFC, 350,000 cells were seeded in 2 mL of culture medium per well in six-well plates containing coverslips (VWR, 16004-326) and incubated at 37 °C and 5% CO2 for 16-24 h, until they reached 60-80% confluence. BiFC assays used a three-plasmid transfection system (VN, CC and mCherry as a transfection control). For each condition, 1 µL of each plasmid solution was added to 200 µL of jetPRIME buffer (VWR, CA89129-922) in a 1.5-mL tube, vortexed and spun down. jetPRIME reagent was added at a 1:2 ratio (µg DNA:µL reagent), and the mix was briefly vortexed, spun down and incubated for 10 min at room temperature. The 200 µL of transfection mix was then added dropwise to a well and distributed evenly. Cells were incubated for 24 h at 37 °C and 5% CO2 before imaging by a Leica SP8 confocal microscope. For each replicate, at least three images were captured from different areas, each containing at least 100 cells. Maximum intensity projections of the z-stacks were quantified with Fiji. For each condition, two fields were analyzed, and in each field the mean mCherry and mean VN/CC (BiFC) fluorescence intensities were measured. The BiFC signal was normalized by dividing the mean VN/CC intensity by the mean mCherry intensity.

### Skeletal preparation

Limbs were dissected from E15.5 embryos and put overnight in ethanol 95%, then stained with Alcian blue for 24h, rinsed with ethanol 95% for 24h, dissolved in KOH 1% for 2 h and stained with Alizarin red for a few hours. Limbs were finally dehydrated in a series of ethanol: glycerol: water solutions and kept in a final solution of ethanol: glycerol (1:1).

### Immunostaining, image acquisition and quantification

Cryosections were made by fixing the E12.5 limb buds overnight in 4% PFA at 4 °C, then put in a 30% Sucrose solution, embedded in 30% Sucrose/OCT (1:1) and sectioned at 12 µm. Sections were blocked 1 h with a saturation solution (BSA 1%, Triton 0.4%) and incubated overnight with antibody, then incubated 45 min with secondary antibody (1:1000) and mounted in a DAPI mounting medium. Sections were imaged with a Leica SP8 confocal microscope.

For whole mount immunofluorescence, collected embryos were fixed overnight in Methanol: DMSO (1:1) at 4 °C, then bleached with H_2_O_2_ in the fixative solution (Methanol: DMSO: H_2_O_2_, ratio 4:1:1) for 5 hours at room temperature, washed with Methanol and then stored at −20 °C. For the staining, forelimbs were dissected and rehydrated in a graded series of Methanol: PBST (PBS, 0.5% Triton), then blocked 2 hours in PBSMT (PBS, 2% silk milk, 0.5% Triton), and incubated overnight at 4 °C with the primary antibody. The next day, forelimbs were washed for a total of 5 hours in PBSMT at room temperature and then incubated with secondary antibody diluted 1:500 in PBSMT overnight at 4 °C. Forelimbs were then washed in PBMST, PBST, and PBS, three times 30 minutes each, then put in a Fructose: Glycerol solution overnight before being imaged with a Leica SP8 confocal microscope.

With iHOXA13 and iHOXA13^F346A^ iPSC, cells adherents on coated VTN-N 12mm coverslip #1 (Fisherbrand, 12541000) cells were fixed for 10 min in 4% PFA at 4 °C, then blocked for 10 min in blocking solution (PBS, 10% FBS, 0.2% Triton) for 10 min. Primary antibodies were diluted in a staining solution (PBS, 2% FBS, 0.2% Triton) and incubated overnight at 4 °C in a humidified chamber. After three PBS washes, secondary antibodies (1:1000) were added for 1h at room temperature. After three PBS washes, cells were then mounted in DAPI mounting medium (Abcam, ab104139). Cells were visualized and imaged using a DM6000 microscope (Leica). For each replicate, at least three images were captured from different areas, and the nuclear signal intensity was then quantified using CellProfiler v4.2.6. Primary antibodies are listed in Supplementary Table 4.

### Quantitative RT-PCR analysis

Total RNA was extracted (Qiagen, 74134) and cDNA synthesized using M-MuLV reverse transcriptase (NEB, M0253). Quantitative real-time PCR was performed using a QuantStudio 5 Real-Time PCR System (Thermofisher Scientific) with PowerUp SYBR Green Master Mix (ThermoFisher, A25741). All expression data were normalized to GAPDH mRNA (ΔCt). All analyses were performed with three technical replicates per plate. mRNA expression levels are represented as relative expression levels determined by calculating 2^−ΔΔCt^. Primers are listed in Supplementary Table 4.

### Ethics statement

The animal study was approved by the Animal Care Committee of IRCM (protocol 2024-03 KM)

## Author contributions

MK and FG designed the study. RR performed all the iPSCs experiments. RPaul performed the *in vivo* experiments. RS carried out the RIME assays and analysis. RPeraldi generated the immunofluorescence on mouse limb buds. MK, RR, FG, SS, RPaul analyzed the data. MK wrote the manuscript with the help of FG, SS and RR. All authors commented on the manuscript.

## Acknowledgements

We thank Sarah Boissel and the members of the Molecular Biology and Functional Genomics Core Facility at the IRCM for generating the sequencing libraries. We also thank the IRCM Imaging Facility for the availability and maintenance of the microscopes, the IRCM Proteomics Facility for the mass spectrometry analyses, and the IRCM Bioinformatics Facility for their advice. We thank Valérie Magoon for her administrative assistance, and we are particularly grateful to the members of the Kmita lab for helpful discussions. Bioinformatics analyses were enabled in part by support provided by Calcul Quebec (www.calculquebec.ca) and the Digital Research Alliance of Canada (www.alliancecan.ca).

