## Supplemental Figures for "HOX13 factors shape cell developmental transcriptional program through multiple mechanisms that regulate chromatin accessibility"

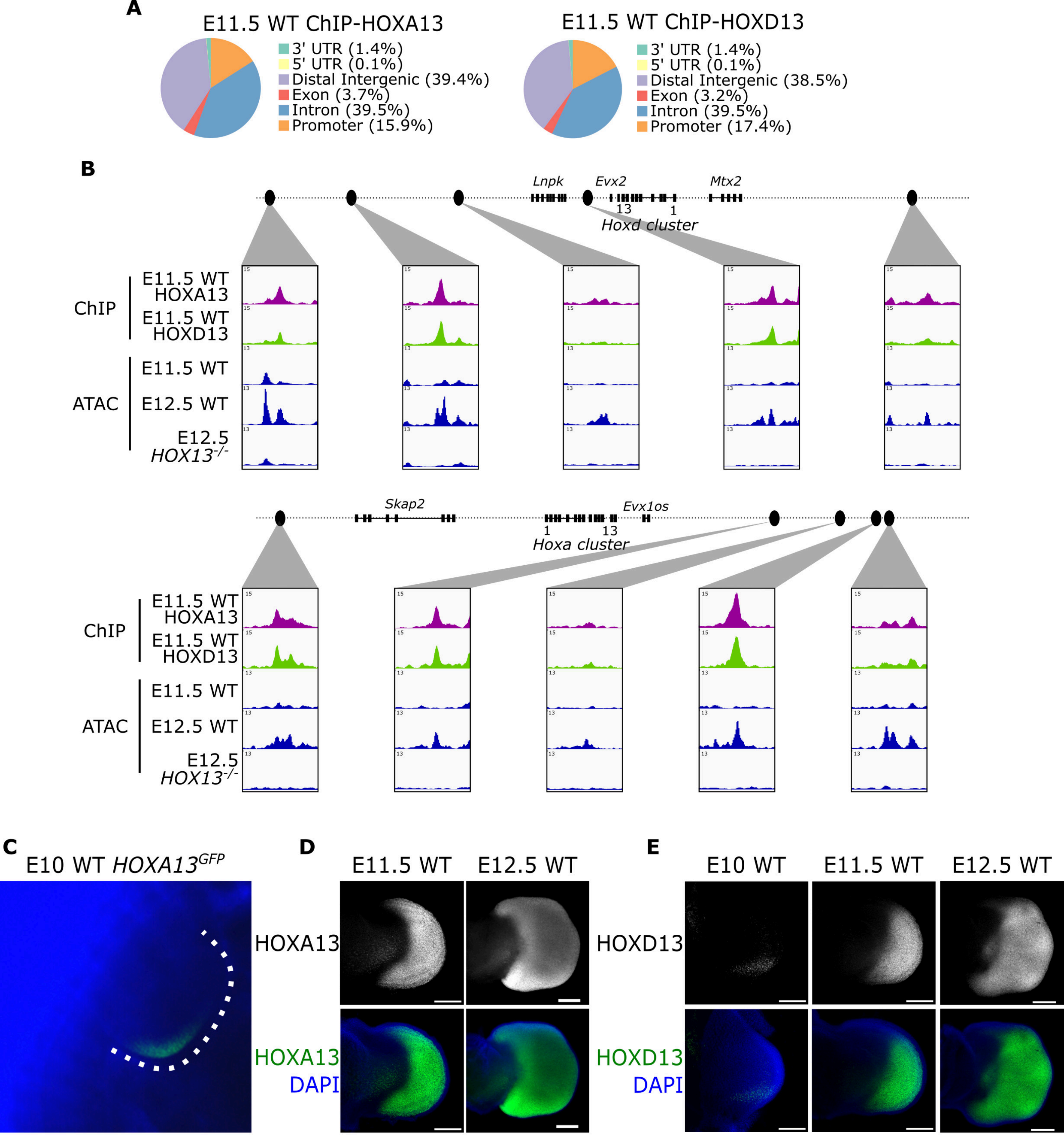


Figure supplementary 1: (**A**) Annotation of E11.5 WT ChiP-seq peaks of Hoxa13 and Hoxd13. (**B**) Genome browser views (IGV) of the Hoxa and Hoxd clusters showing ChIP-seq and ATAC-seq signal in the mouse distal limb. (**C**) Expression of the Hoxa13-GFP reporter in a heterozygous E10.0 forelimb. (**D**) Whole-mount immunostaining for HOXA13 in wild-type mouse limbs at E11.5-E12.5. (**E**) Whole-mount immunostaining for HOXD13 in wild-type mouse limbs at E10.0-E12.5. Scale bars, 200 μm (**D**), 200 μm (**E**).

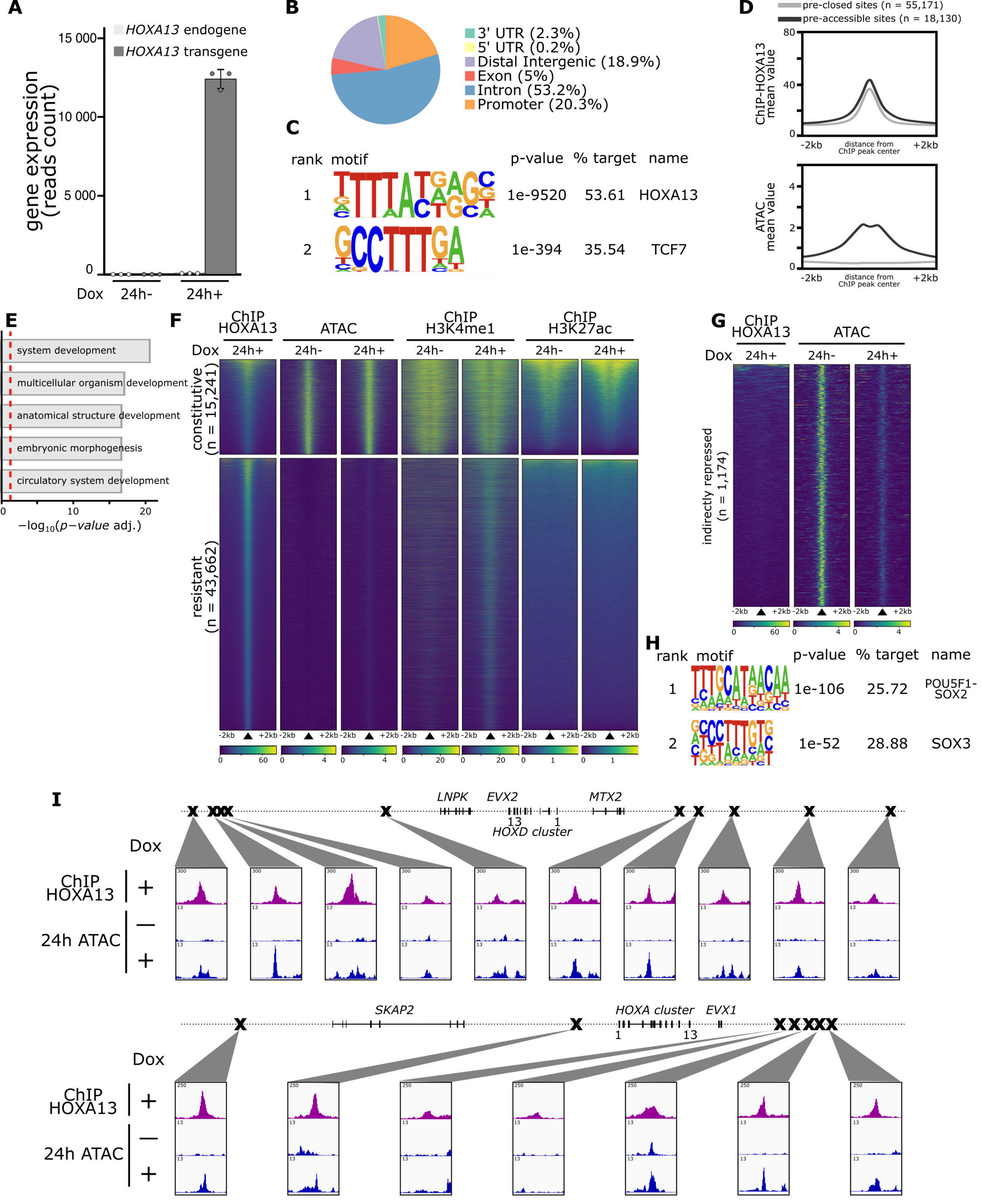


Figure supplementary 2: (**A**) Gene expression (RNA-seq) of the endogenous and transgene HOXA13 transcripts detected in iHOXA13-iPSCs with or without Dox stimulation. (**B**) Annotation of the HOXA13 ChIP-seq peaks (n = 73,301) in iHOXA13-iPSCs. (**C**) Top-scoring de novo motifs identified by HOMER in a 200 bp window centered on the HOXA13 binding sites (n = 73,301). (**D**) Average profiles of HOXA13 ChIP-seq and ATAC-seq signal at sites closed before HOXA13 recruitment (n = 55,171) or already accessible (n = 18,130). (**E**) GREAT analysis of enriched Gene Ontology (GO) biological process terms for genes associated with the HOXA13 binding sites (n = 73,301). (**F**) Heatmaps of the constitutive (n = 15,241) and resistant (n = 43,662) HOXA13 sites, showing HOXA13 ChIP-seq signal at 24 h of Dox treatment (n = 2), and H3K4me1 and H3K27ac ChIP-seq and ATAC-seq signal at 24 h without treatment (n = 2) or with Dox (n = 2) in iHOXA13-iPSCs. Sites are ranked by decreasing HOXA13 ChIP-seq signal. (**G**) Heatmaps of the sites closed indirectly through HOXA13 induction (n = 1,174), showing HOXA13 ChIP-seq signal at 24 h of Dox treatment (n = 2) and ATAC-seq signal at 24 h without treatment (n = 2) or with Dox (n = 2) in iHOXA13-iPSCs. Sites are ranked by decreasing HOXA13 ChIP-seq signal. (**H**) Top-scoring de novo motifs identified by HOMER in a 200 bp window centered on the sites closed indirectly through HOXA13 induction (n = 1,174). (**I**) Genome browser views (IGV) of the HOXA and HOXD clusters showing ChIP-seq and ATAC-seq signal in iHOXA13-iPSCs.

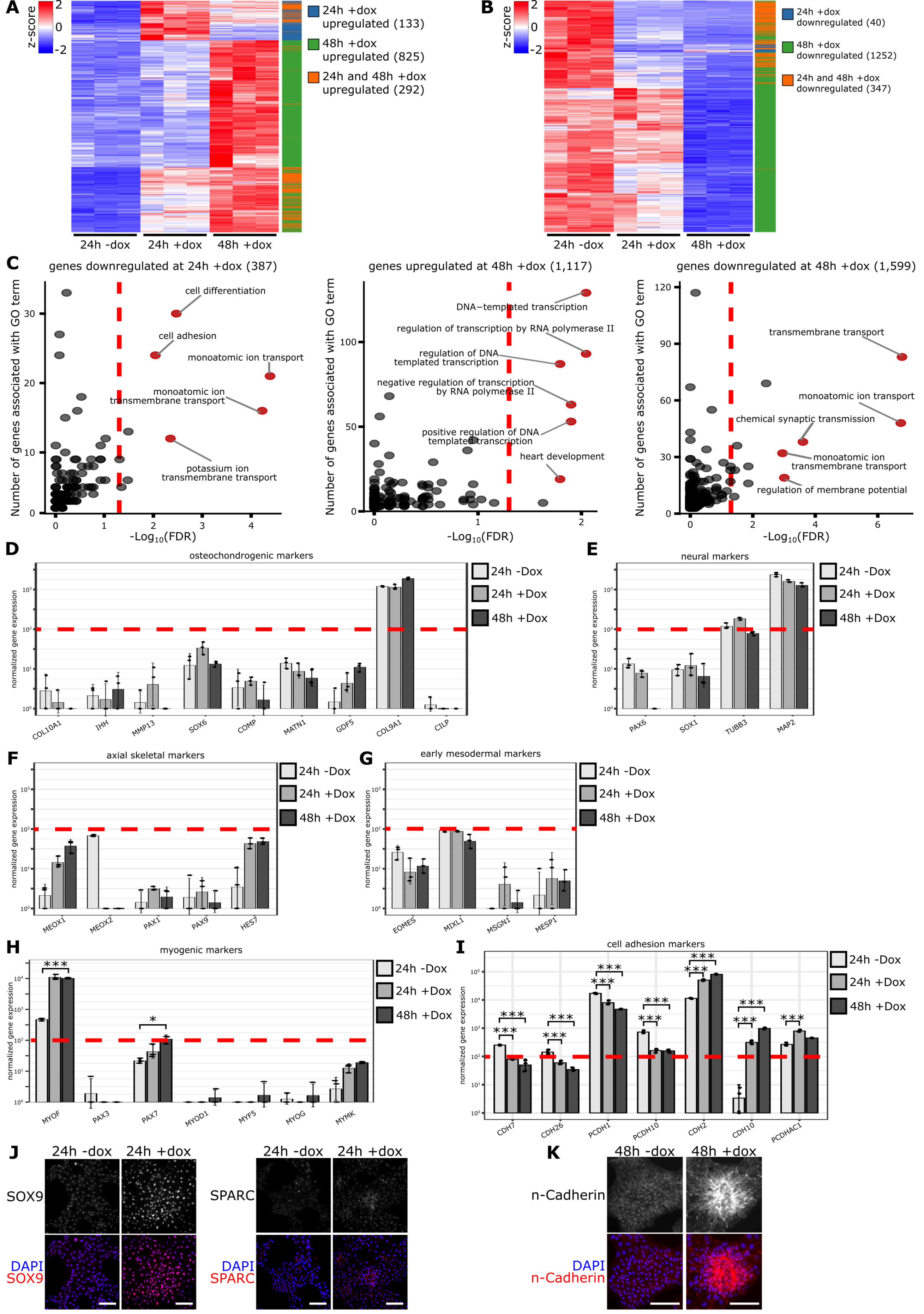


Figure supplementary 3: (**A-B**) Hierarchical clustering showing transcriptional dynamic (z-score) of upregulated genes (**A**) or downregulated genes (**B**) after 24 h and 48 h Dox treatment. (**C**) Functional enrichment analysis (Gene Ontology: Biological Process) of the 387 genes downregulated at 24h + Dox (left), 1,117 genes upregulated at 48h + Dox (middle) and 1,599 genes downregulated at 48h + Dox (right). The y axis shows statistical significance ($-\log_{10}$ false discovery rate, FDR) and the x-axis represents the number of genes associated with each GO term. (**D-I**) Barplots of RNA-seq normalized gene expression of osteochondrogenic (**D**), neural (**E**), axial skeletal (**F**), early mesodermal (**G**), myogenic (**H**) and cell adhesion (**I**) markers. Red dashed lines indicate the cutoff (≥ 100) used to distinguish expressed from non-expressed genes. (**J**) Immunostaining for SOX9 (osteochondrogenic marker) and SPARC (osteogenic mesenchymal marker) in plated iHOXA13-iPSCs treated or not with Dox during 24 h. (**K**) Immunostaining for n-Cadherin in plated iHOXA13-iPSCs treated or not with Dox during 24 h. Scale bars, 100 µm (**J**), 100 µm (**K**).

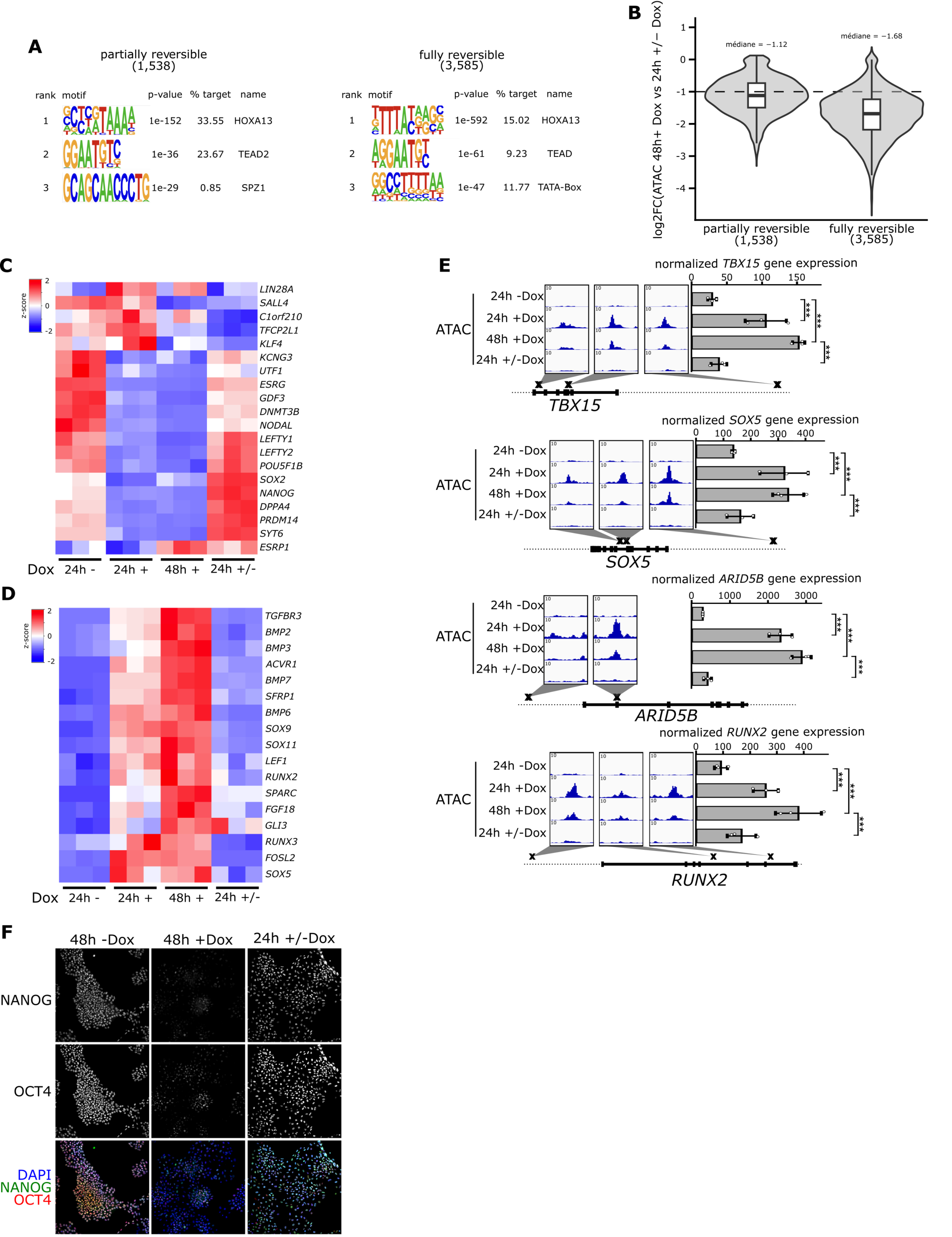


Figure supplementary 4: (**A**) Top scoring de novo motifs identified by HOMER in a 200 bp window centered on sites that decreased in accessibility (n = 1,538) or completely lost accessibility (n = 3,585) after Dox withdrawal. (**B**) Violin plots with overlaid box plots showing the log2 fold change in ATAC-seq signal (48h + Dox vs 24h +/− Dox) at the sites with decreased in accessibility (n = 1,538) or completely lost accessibility (n = 3,585) after Dox withdrawal. (**C**) Heatmap of RNA-seq gene expression (row z-score) for pluripotency-associated genes after Dox withdrawal. (**D**) Heatmap of RNA-seq gene expression (row z-score) for osteoblast and chondrocyte-associated genes upregulated after 24 h and/or 48 h of Dox treatment, shown after Dox withdrawal. (**E**) Genome browser views (IGV) of *TBX15*, *SOX5*, *ARID5B* and *RUNX2* loci, showing ATAC-seq and RNA-seq signals in 24 h - Dox, 24 h + Dox, 48 h + Dox and 24 h ± Dox. (**F**) Immunostaining for NANOG and OCT4 in iHOXA13-iPSCs treated with or without Dox for 48 h, and after Dox withdrawal

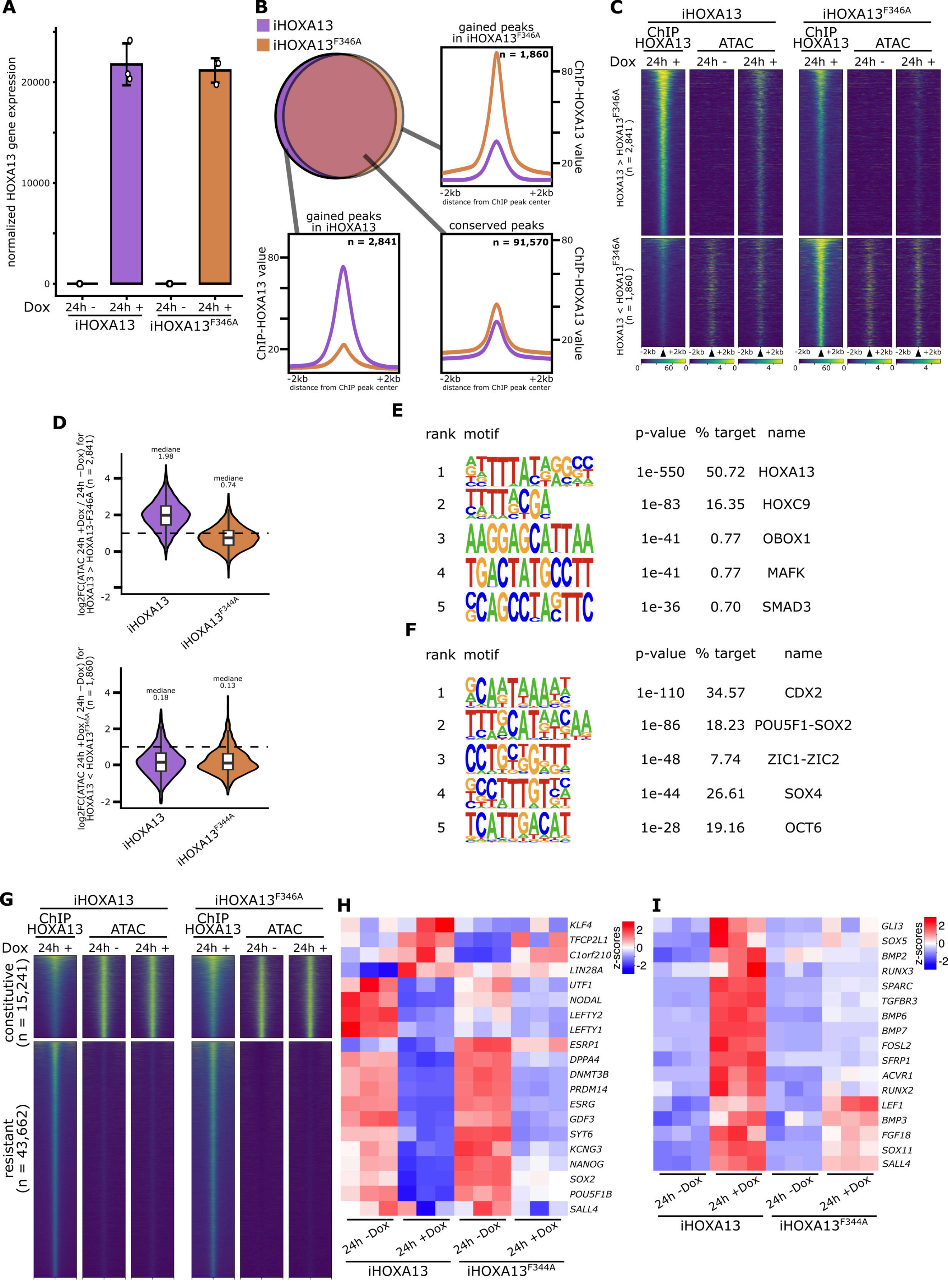


Figure supplementary 5: (**A**) Barplots of RNA-seq normalized gene expression of HOXA13 transgen in iHOXA13 and iHOXA13$^{F346A}$-iPSC. (**B**)Comparison and average profils of ChIP-seq peaks in iHOXA13 (purple) and iHOXA13$^{F346A}$-iPSCs (orange). (**C**) Heatmaps of positions with higher binding of HOXA13 (n = 2,841) and HOXA13$^{F346A}$ (n = 1,860). Sites are ranked by decreasing HOXA13 ChIP-seq signal. (**D**) Violin plots with overlaid box plots showing the $\log_2$ fold change in ATAC-seq signal (24 h +Dox vs. 24 h −Dox) at the sites with higher binding of HOXA13 (n = 2,841) and higher binding of HOXA13$^{F346A}$ (n = 1,860), in iHOXA13 (purple) and iHOXA13$^{F346A}$-iPSCs (orange). (**E-F**) Top scoring de novo motifs identified by HOMER in a 200 bp window centered on sites with higher binding of HOXA13 (n = 2,841) (**E**) and higher binding of HOXA13$^{F346A}$ (n = 1,860) (**F**). (**G**) Heatmaps of the constitutive (n = 15,241) and resistant (n = 43,662) HOXA13 sites, showing HOXA13 ChIP-seq signal after 24 h of Dox and ATAC-seq signal after 24 h without treatment or with Dox in iHOXA13 and iHOXA13$^{F346A}$-iPSCs (n = 2 for each ChIP-seq and ATAC-seq condition). Sites are ranked by decreasing HOXA13 ChIP-seq signal. (**H-I**) Heatmap of RNA-seq gene expression (z-score) for pluripotency (**H**) and osteoblast/chondrocyte-associated (**I**) genes in iHOXA13 and iHOXA13$^{F346A}$-iPSCs with or without Dox (n = 3 for each RNA-seq condition)

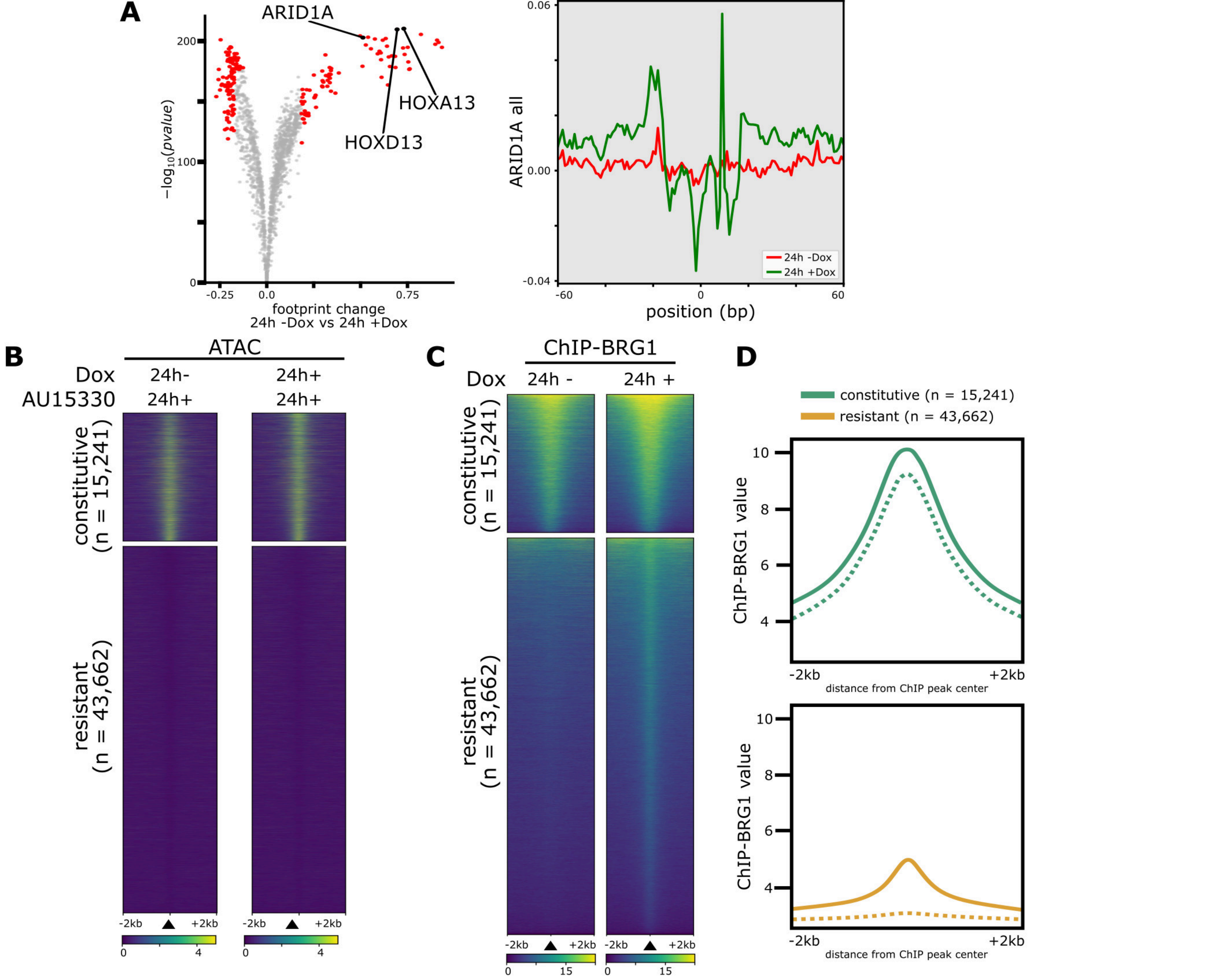


Figure supplementary 6: (**A**) Differential footprint analysis of ATAC-seq signal at 24 h without treatment (n = 2) or with Dox (n = 2) in iHOXA13-iPSCs (left) and aggregate footprint profiles at ARID1A motifs across all sites with gained accessibility (right). (**B**) Heatmaps of the constitutive (n = 15,241) and resistant (n = 43,662) HOXA13 sites, showing BRG1 ChIP-seq signal in iHOXA13-iPSC treated with AU-15330 for 24 h, without Dox (n = 2) or with Dox (n = 2). Sites are ranked by decreasing HOXA13 ChIP-seq signal. (**C-D**) Heatmaps (**C**) and average signals (**D**) of the constitutive (n = 15,241) and resistant (n = 43,662) HOXA13 sites, showing BRG1 ChIP-seq signal after 24 h without treatment (n = 2) or with Dox (n = 2) in iHOXA13. Sites are ranked by decreasing HOXA13 ChIP-seq signal.
